# Lifespan Trajectories of Resting State Bipolar EEG Power

**DOI:** 10.64898/2026.09.28.755104

**Authors:** David B. Chorlian, Chella Kamarajan, Jacquelyn L. Meyers, Andrey Anokhin, Ashwini K. Pandey, Jian Zhang, Sivan Kinreich, Bernice Porjesz

## Abstract

Normative trajectories of resting state bipolar EEG power over the lifespan were derived from a sample of 5781 participants from the Collaborative Study on the Genetics of Alcoholism (COGA), a large multi site study, using 12,906 observations from ages 12 through 70. Trajectories were analyzed using the means, variances, and correlations of data in seven frequency bands (from low theta to high beta) and 24 bipolar channels obtained from 17 electrodes of the 10-20 system. In order to make the study more useful for understanding EEG in the general population, only subjects never having an alcohol use disorder (AUD) diagnosis were included in the study. Bipolar EEG data is derived from the differencing of the signals in adjacent electrodes, either in a lateral (left-right) or sagittal (anterior-posterior) orientation; each difference is called a channel. The power values for the individual observations were calculated by standard Fourier transform based methods.

In all cases there are sharp decreases in EEG power from ages 12 to the early 20s after which there are number of divergent patterns: general increases in the beta frequencies, and mixed patterns of increase and subsequent decrease in the theta and alpha frequencies. In these characteristics the lifespan trajectories presented here are similar to the monopolar trajectories described in our previous study Chorlian et al. (2026). Sagittal and lateral bipolar trajectories are similar within each sex and between sexes; the primary difference is the much lower degree of regional difference in the sagittal trajectories as compared to the lateral trajectories. The pairwise inter-frequency (within channel) correlation trajectories between the four theta and alpha frequency bands and the four high alpha and beta frequency bands at each channel show striking sex differences unlike the corresponding monopolar correlations. There are a number of features in the bipolar power trajectories that suggest that the differencing procedure reduces not only local phase synchrony, but also effects of global and regional phase synchrony present in the monopolar data from which the bipolar signals are derived. The inter-frequency correlations at the individual channels are much higher in the bipolar data than the monopolar data, suggesting that global and regional phase synchrony components retained in the monopolar data have greater frequency specificity than the more purely local components in the bipolar data. Further analyses to examine many other features of the data will be undertaken in our subsequent work.

## 1 Introduction

The Collaborative Study on the Genetics of Alcoholism (COGA) (Begleiter et al., 1995; Agrawal et al., 2023; Meyers et al., 2023) has published many studies of resting state EEG since its inception covering both power and coherence primarily related to alcohol use disorder related conditions (Rangaswamy et al., 2002, 2003), but also genetic aspects (Porjesz et al., 2002; Tang et al., 2007a,b; Chorlian et al., 2007), and combinations of these factors (Meyers et al., 2017, 2019, 2021a,b; Chorlian et al., 2023). COGA is a participant in the ENIGMA-EEG working group and involved in projects described in Smit et al. (2018) and Smit et al. (2021). In our recent study of resting state EEG monopolar power values we reported estimations of mean values of power and of both their inter-frequency and intra-frequency correlations extending from age 12 to age 70 (Chorlian et al., 2026). This study of bipolar data is a complement to our previous study; the rationale for the use of bipolar data is twofold: it is independent of reference and it removes some volume conduction effects (Nunez and Srinivasan,, 2006; Chorlian et al., 2009), particularly important in studies using relatively few electrodes. EEG bipolar data should be regarded as a combination of the unique features of the data at each of the electrodes whose difference is calculated together with the removal of the features common to the electrodes, whether these features are common only to that pair or extend over the entire set of electrodes in the study. Using the same methodology of the monopolar study with a somewhat larger number of participants, we performed a comprehensive analysis of resting state EEG bipolar power values extending from age 12 to age 70, which included estimations of mean values and of both inter-frequency and intra-frequency correlations. Specifically, standard Fourier transform methods were used to estimate power in 1/2 Hz bins and aggregated in seven frequency bands used in previous studies. This data was then subject to a non-parametric regression analysis and complementary covariance estimation at 117 age centers 1/2 year apart from age 12 to age 70. Additional calculations were made in order to quantify male/female differences and other features of the power and correlation trajectories. While this study was initially intended to be completely self-contained study without reference to the monopolar results, we found it provided considerable additional insight to the interpretation to contrast the two sets of results even without explicit quantification of their relations, which we will consider in a subsequent study.

We observed a variety of features which we believe are previously unreported in the EEG literature. We note that our observations are not uniformly distributed across the analyzed age range, being highly skewed toward the younger ages; we thought it desirable to present these because the effective sample sizes were large compared to many EEG studies, as explained in section 2.3.1. We used conventional frequency band based analysis in order to make possible comparisons to earlier studies, particularly our monopolar power study. This is the second of series of studies using an identical methodology; subsequent studies will cover bipolar EEG coherence and a comparison of the monopolar and bipolar power results using methods additional to those used here. In order for this document to be self-contained as possible some sections of this study are copied directly from our previous study (Chorlian et al., 2026).

### Authors’ note

We urge readers to open two instances of their PDF viewer in order to more easily compare the descriptions of the results with the figures illustrating them as these are in separate sections of the document. We also suggest the readers take the opportunity to increase the magnification of the figures to check the details of the description when desired, as the plots provided will produce precise results when magnified by factor of 4 at least.

## 2 Materials and Methods

### 2.1 Sample

As part of the Collaborative Study on the Genetics of Alcoholism (COGA) (Begleiter et al., 1995; Agrawal et al., 2023), EEG data were collected from families densely affected with alcohol use disorder (AUD) and their offspring as well as community comparison families. Many of the initially recruited subjects were recontacted recently and additional EEG data was collected. This provides a sample of subjects with observations ranging from age 10 to over 70 years of age. A phase of the study in which the offspring of previous participants were tested repeatedly from ages 12 and onward at approximately 2 year intervals provided a large number of observations from individuals under age 30. In order to make this study more comparable to results from the general population, all data from subjects who at any age had a DSM-5 AUD diagnosis were removed from the study. Subjects were not excluded for other alcohol use behavior.

### 2.2 Data Collection and Processing

Data was collected as described in Chorlian et al. (2009) and Meyers et al. (2019). Subjects were asked to keep their eyes closed and not to move their head while data from 19 or more electrodes of the 10-20 system was recorded for a period of 256 seconds in an enclosed booth. In the interest of retaining data from as many subjects as possible, we used 17 of the basic 19 electrodes of the 10-20 system for our analysis. Data cleaning was effected by a subspace removal method, removing a portion of the data from the scalp channels correlated to the eye channel. The one second segments in which the maximal difference among observations at any one electrode exceeded 100 micro volts were marked as unusable. Subject data without at least 129 usable segments was rejected. For observations with usable data, the 24 bipolar channels were created by the differencing of pairs of electrodes as indicated in section 5.3. After the differencing, the data was rechecked for usability in the same manner as that used for the original data. The 24 channels created in the differencing procedure fall in two groups of 12 each, depending on the orientation of the pairs of electrodes differenced. The lateral group of channels falls into 3 subgroups of equal size, frontal, central, and parietal. The sagittal group of channels falls into 3 subgroups of sizes 5, 5, and 2, frontal, central, and posterior. The 2 occipital electrodes occur only in the sagittal group; each of the other 15 electrodes appear in both of the groups. The sagittal power will exhibit consistently less regional variation than the lateral power because data from the central electrodes is represented in both the frontal and central regions of the sagittal channels; there is no similar duplication in the lateral channels. See figure 1 for an illustration of the channels.

**Figure 1:**
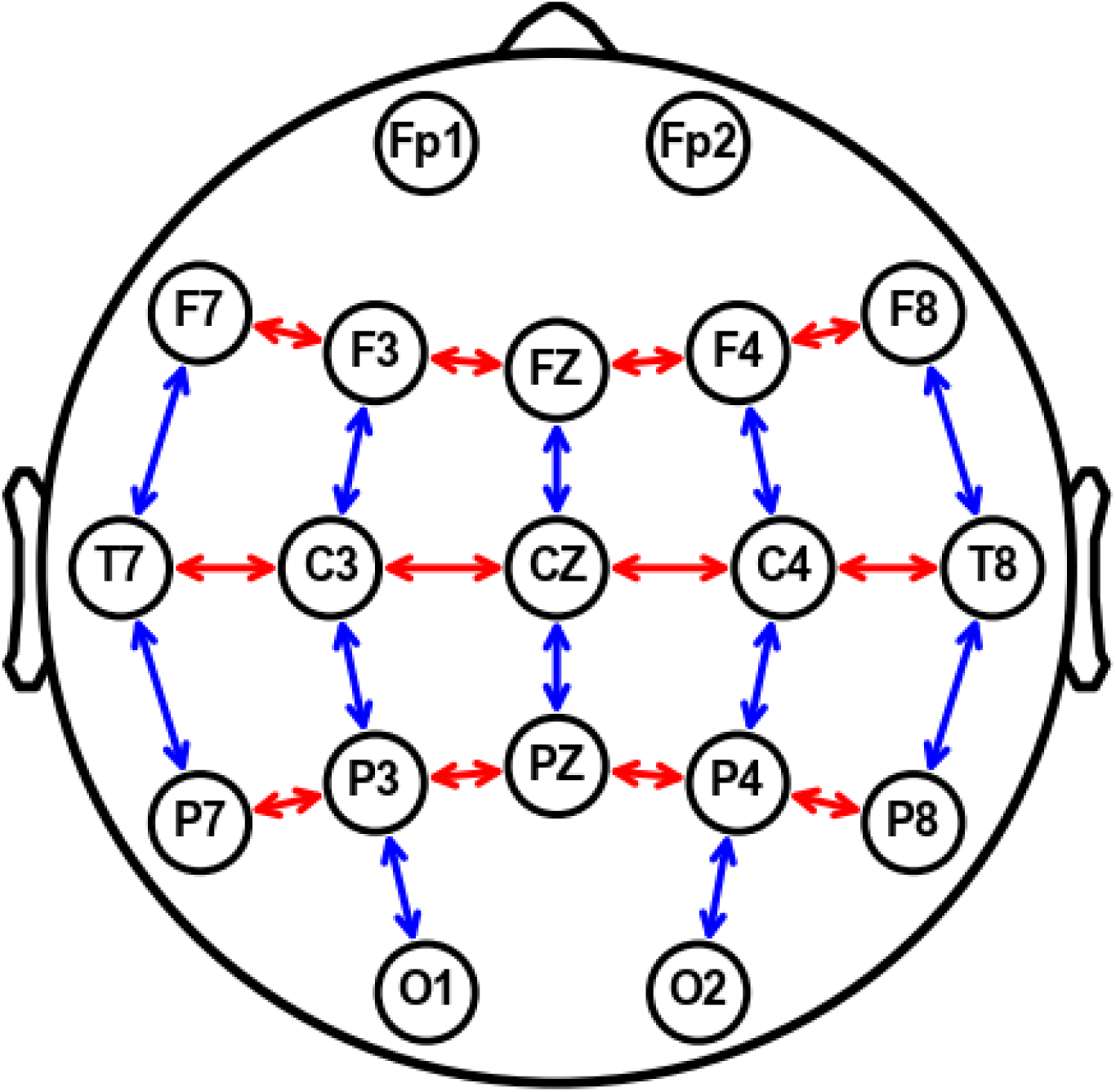
Template of latteral and sagittal pairs. Blue arrows connect sagittal pairs. Red arrows connect lateral pairs.

Power was calculated using a variation of the method used in Smit et al. (2018). Cleaned data were imported into MATLAB, organized into overlapping 2 s epochs, detrended, and power spectra calculated using the Fast Fourier Transformation (FFT). Power was defined as the squared radius of the orthogonal sine and cosine amplitudes averaged over the window size, and the mean value taken for the frequency band to obtain power density. Power values were aggregated into seven frequency bands from low theta to high beta specifically, low theta (3.0-5.0 Hz), high theta (5.0-7.0 Hz), low alpha (7.0-9.0 Hz), high alpha (9.0-12.0 Hz), low beta (12.0-16.0 Hz) mid beta (16.0-20.0 Hz), high beta (20.0-28.0 Hz), and log transformed to obtain approximate normality. Details of the frequency bands and the electrodes used may be found in section 5. Power as calculated from the difference of two electrodes is a function of the power at the individual electrodes and of the phase relations between the signals. That is, there is no direct relation between the bipolar power calculated here and the monopolar power of the individual electrodes in the differencing. The amount of phase synchronization across the scalp affects the power estimation. When signals exhibit phase synchronization, the distance between any two signals is bounded by the difference between the largest amplitude and the smallest amplitude; in the absence of synchronization the distance is bounded by the sum of the distances of the two largest signals. This leads to the possibility of the reversal of power relations between the monopolar data and the bipolar data. No attempt was made to measure the amount of phase synchronization of the data.

### 2.3 Analysis Methods

#### 2.3.1 Regression Modeling

A non-parametric local linear regression (LOESS) method was used to estimate the mean and variance of the power data at 117 age centers 1/2 year apart for ages from 12 to 70. The bandwidth for the kernel (Epanechnikov) was taken as 0.6. More specifically, the model for each age *t_k_, k* = 1 *. . .* 117, is represented by the following equation, in which *y_i j_* is the dependent variable for subject *i* at observation *j* at age *t_i j_*, *W_h_*(*t_i j_* − *t_k_*) the kernel weighting for the local linear regression model, a function of the bandwidth (fraction of data included) *h*, *β*_0_*_,n,k_* the parameter to be estimated for covariate *n* for its effect on the intercept and *β*_1_*_,n,k_* the parameter to be estimated for covariate *n* for its effect on the slope, and *X_i, j,m_* the value of covariate *m* of subject *i* at observation *j*. Different sets of covariates may be used for the estimation of the slope and the intercept. The weight *W_h_*(*t_i j_* − *t_k_*) will be non-zero only for a set of values near *t_k_*. Note that the beta values are indexed to emphasize that each is the product of a separate estimation for each age. (In the equation, intercept terms appear on the first line and slope terms on the second line. The summations are over the repeated indices in the covariates and betas.)

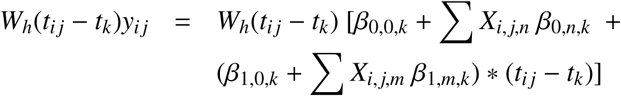

In the statistical model sex is the only covariate for the intercept and slope. Weights for individuals were adjusted to account for multiple observations on single individuals. Results are not independent between models for different ages since the data in each of the 117 regression models has considerable overlap with the data used in the models for nearby ages. This model was first fully described in Chorlian et al. (2015). Unlike in its previous uses, the age distribution of the sample is skewed, with about 3/4 of the observations from subjects thirty years of age or younger. Although 60% of the observations are used in each regression calculation, the values of the weights depends on the distribution of the differences in age from the age center of the observations, and thus the skewed age distribution affects the age distribution of the effective sample size, defined as (*W_h_*(*t_i j_* − *t_k_*))^2^*/* ^.^(*W_h_*(*t_i j_* − *t_k_*)^2^) for each *t_k_*, as illustrated for this data set in figure 6.

**Figure 2:**
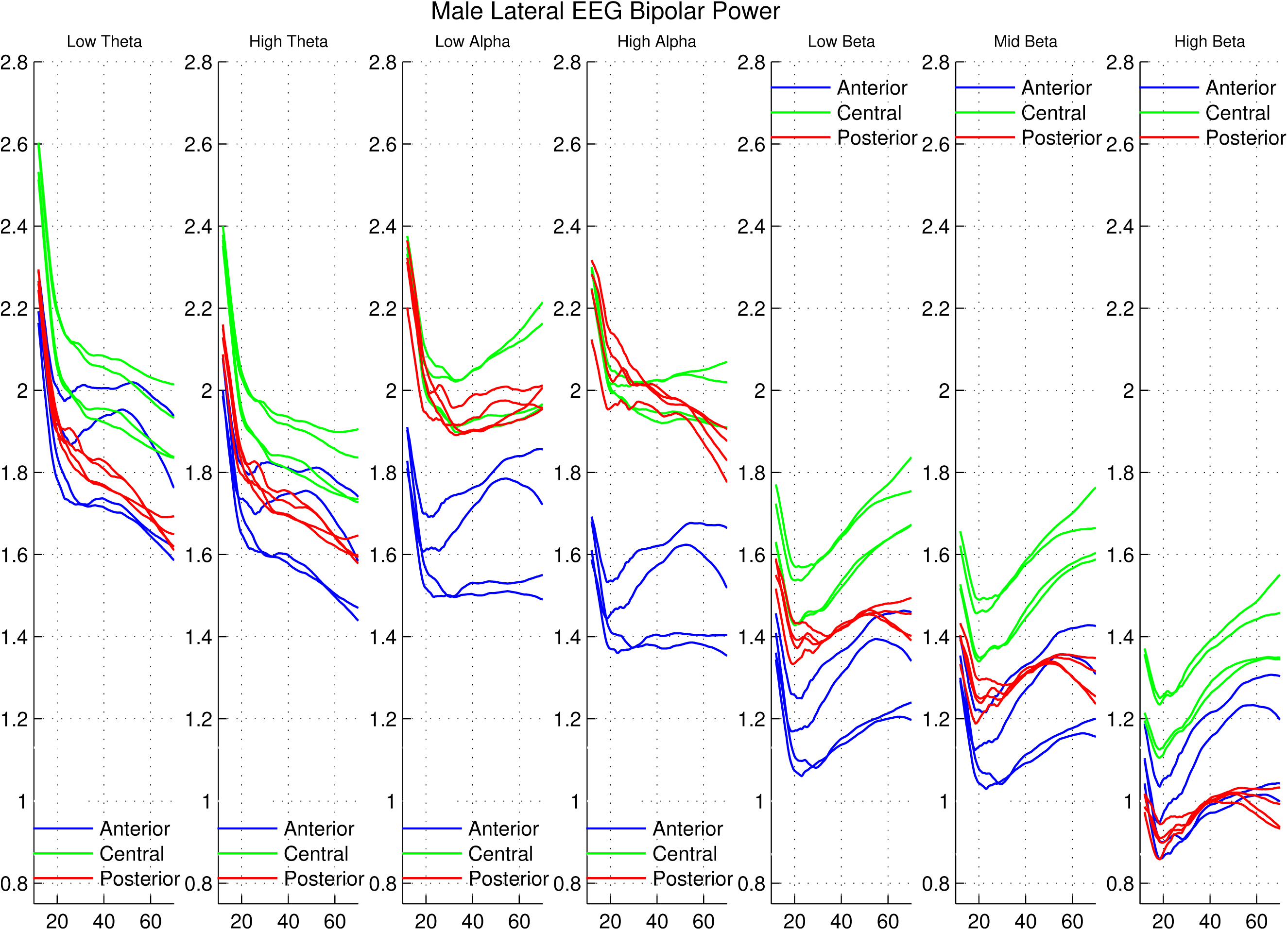
Trajectories of the seven frequency band lateral male means color-coded by region. Anterior is red, central green, posterior red. There are 4 channels in each region.

**Figure 3:**
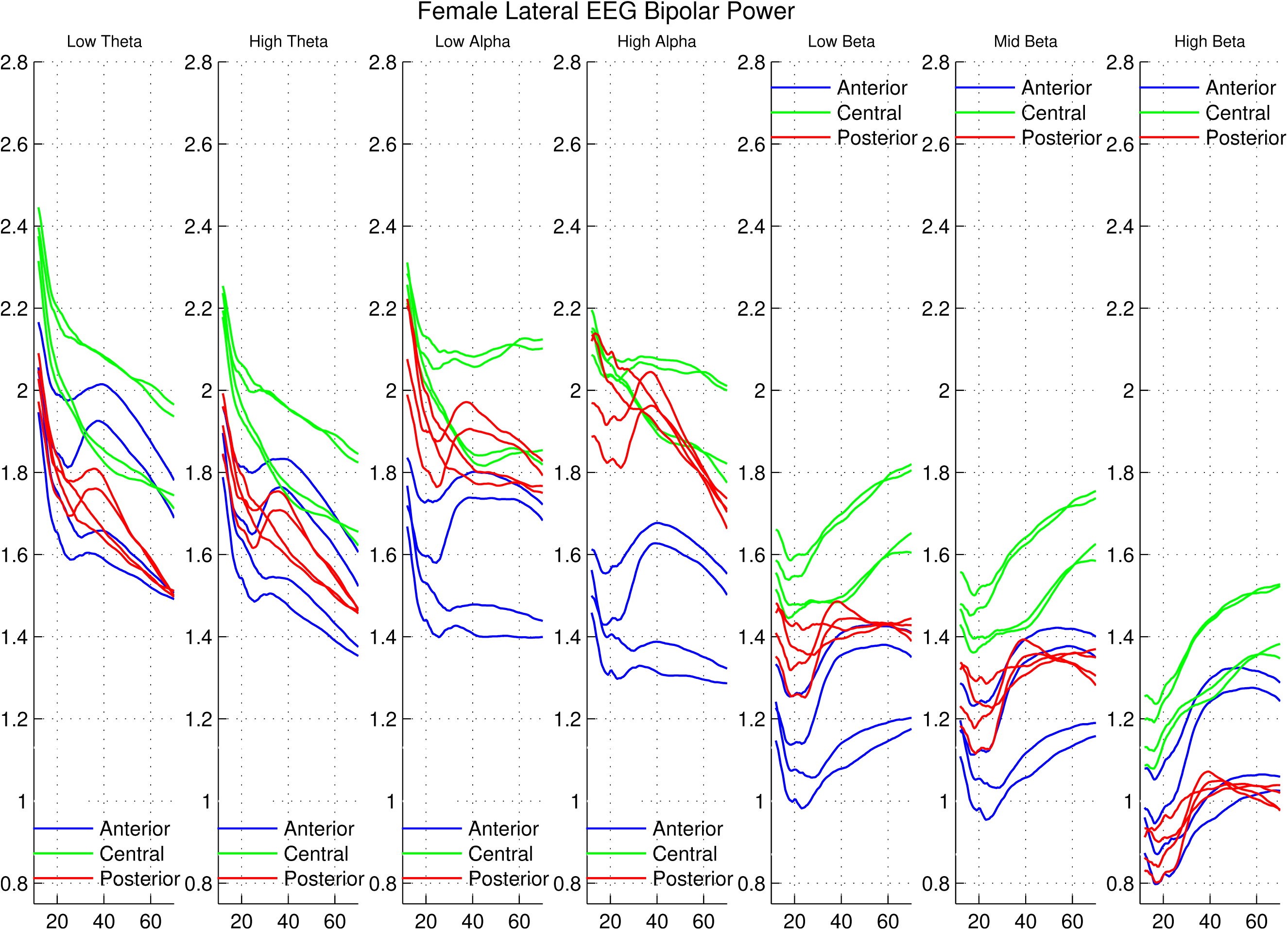
Trajectories of the seven frequency band lateral female means color-coded by region.

**Figure 4:**
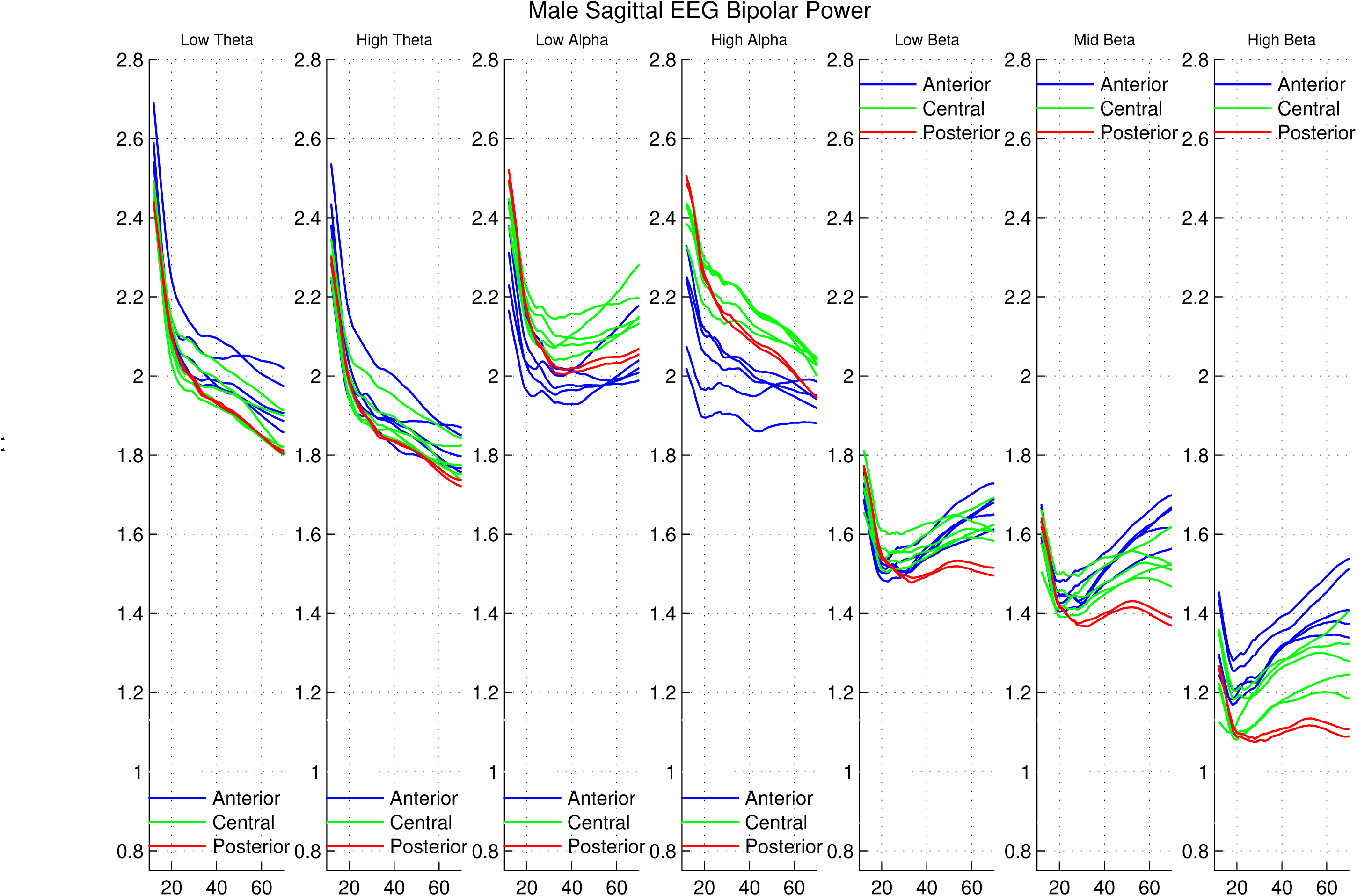
Trajectories of the seven frequency band sagittal male means color-coded by region. Anterior is red, central green, posterior red. There are 5 channels in each of the anterior and central regions, 2 in the posterior region.

**Figure 5:**
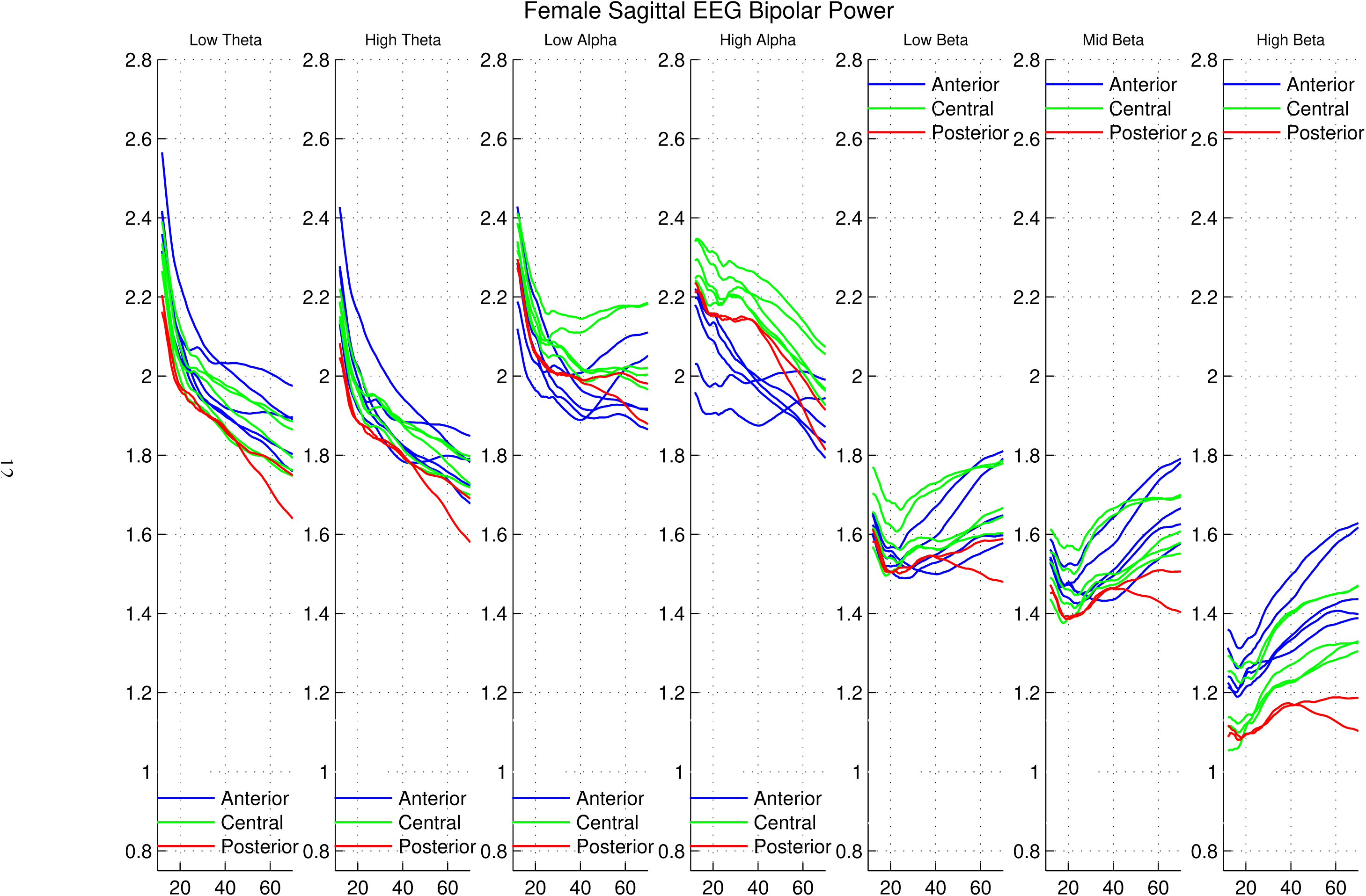
Trajectories of the seven frequency band sagittal female means color-coded by region.

**Figure 6:**
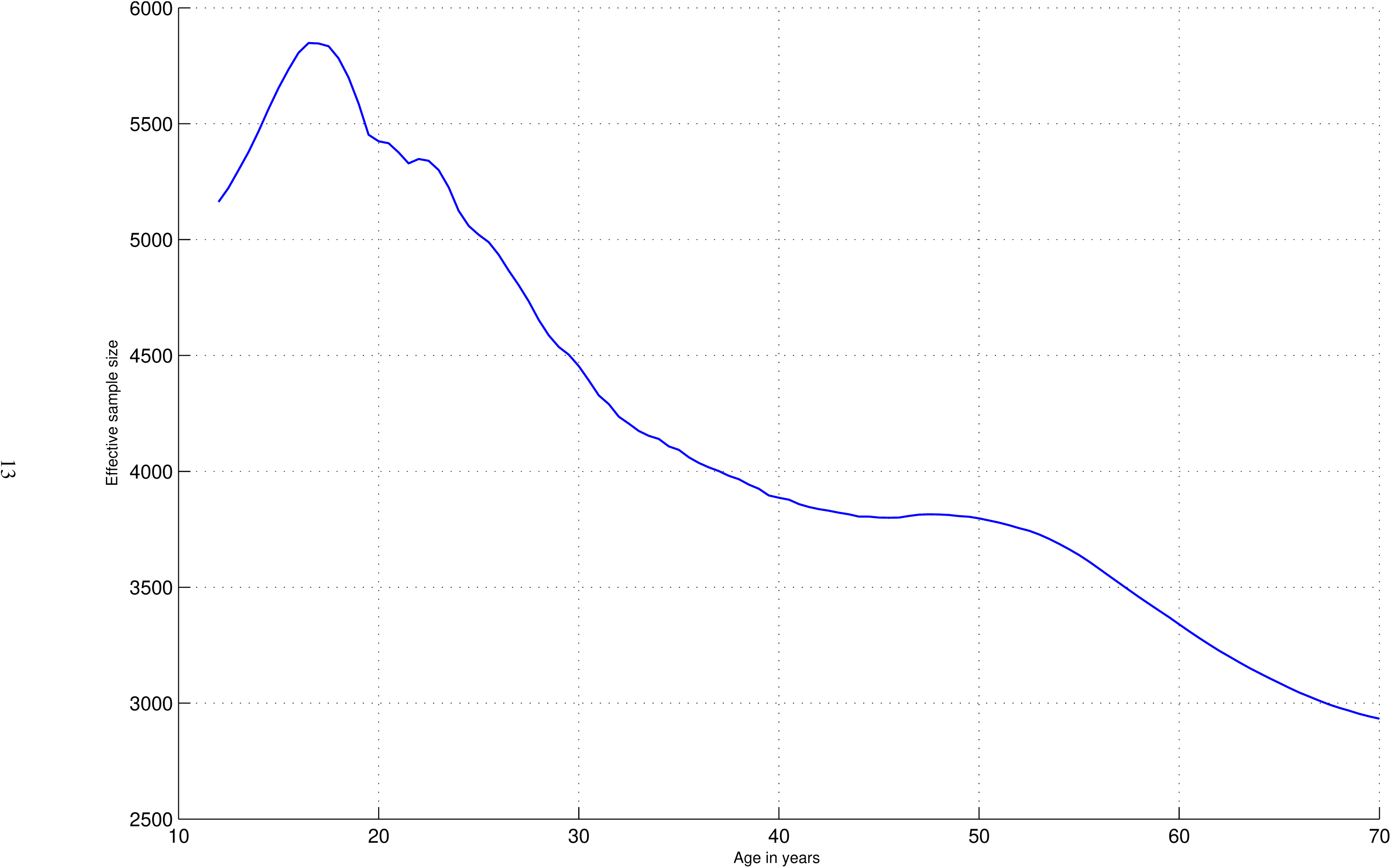
Estimated sample size (bandwidth = 0.6) for regression values.

#### 2.3.2 Covariance Modeling

An adaptation of the regression method, initially described in Chorlian et al. (2015), was used calculate covariances. Briefly, the weights used in the regression calculation were employed in a weighted covariance analysis. The covariance of the data both between all pairs of channels for each frequency band and between all pairs of frequencies for each channel was calculated for each of the age centers. In the covariance analysis males and females were analyzed separately, in contrast to the mean and variance analysis which uses covariates to estimate male and female trajectories. The use of covariates to estimate sex effects provides more accurate estimates of sex differences than separate analyses. Correlational estimates derived from the covariance matrices rather than the covariances matrices themselves were analyzed in order to more clearly differentiate changes in power relations from changes in power values. Although some left/right differences were observed, there is no formal left/right analysis presented because of the greater variation along the anterior/posterior axis, with a secondary variation between peripheral channels and interior channels being clearly visible.

#### 2.3.3 Matrix distance Estimation

Although visual inspection shows many clear differences and similarities of trajectories, we thought it desirable to quantify the difference between comparable sets of trajectories, particularly to characterize male/female differences. To do this, we use the relative Frobenius distance (RFD) between two matrices of identical size and comparable entries. The RFD is the Frobenius norm of the difference of the matrices divided by the mean of Frobenius norms of the individual matrices. This value represents the difference between the matrices in terms of the size of the matrices. It is a dimensionless value. It is not entirely satisfactory when the Frobenius norms of the two matrices differ considerably in size, as the distance between the matrices will be a small fraction of the size of the larger matrix but a large fraction of the size of the smaller matrix. Consequently the only comparisons offered here will be for matrices whose norms are approximately the same size. For example, this rules out direct comparisons between the power trajectories of the beta frequency bands with the theta and alpha frequency bands.

This methodology was applied to quantifying the male/female differences in the power trajectories and in the interfrequency correlation trajectories. For the power case the calculation for each of the 7 frequency bands was made separately for the sagittal and the lateral trajectories. For the correlation cases the calculation was made separately in each of the six theta-alpha correlations and in each of the six high alpha-beta correlations separately for the sagittal and the lateral trajectories. This methodology does not enable the direct comparison of lateral/sagittal differences for although each has 12 channels, the channels do not have topographically similar organizations.

The similarity of power trajectories as measured by RFD may be unrelated to the correlations between the corresponding measures. That is, two measures may have similar trajectories but not be correlated, or two measures may be highly correlated and their trajectories be quite different in shape, as is in fact the case for a number of the measures.

## 3 Results

### 3.1 Introduction

As mentioned previously, EEG bipolar data should be regarded as a combination of the unique features of the data at each of the electrodes whose difference is calculated together with the reduction of the features common to the electrodes, whether these features are common only to that pair or extend over the entire set of electrodes in the study. The consequence of this is that proportionally the more the power at individual electrodes is the result of large scale phase synchronized activity, the less the power of the bipolar channels will be; similarly increases of the power of bipolar channels could be the result of diminished global synchronization. Since we have no independent measures of phase synchrony at this point, we can only suggest its presence in the relation between local and global commonality in the bipolar data. The power calculated from the data is independent of the sign of the data, so the manner in which the differences are calculated is not significant to the results. The sagittal power exhibits consistently less regional variation than the lateral power because data from the central electrodes is represented in both the frontal and central regions of the sagittal channels; there is no similar duplication in the lateral channels. This difference in regional variation is also present in the correlation trajectories.

There are three distinct sets of measures in this analysis. The primary set comprises the direct results of the regression analysis of the 24 channels described above. There are two distinct correlation results, one of the inter-frequency correlations of which we have selected 12 of of the 21 possible pairings of the frequency bands of the 24 channels and the other of the inter-channel (intra-frequency) correlations of which we have selected 23 of the 276 possible pairings. Since the frequency band boundaries are based on aggregated data from previous studies, not on the analysis of the frequential fine structure of the data, examining the inter-frequency correlation structure provides additional information about large scale frequency relations. The inter-channel correlations were examined in order to characterize their consistency across frequency bands. Although plots show that trajectories of inter-channel correlations are similar across frequencies for the inter-channel pairs we examined, we postpone the discussion of these results because our methods of quantifying the similarity across frequency bands are not completely satisfactory.

Results are derived from visual inspection of the trajectories found in the plots found in section 6 and additional quantification of their characteristics described above.

### 3.2 Trajectories of EEG power across the Lifespan

On first inspection of figures 2 through 5 the overall characteristics are the separation of trajectories by frequency type (i.e theta, alpha, beta) in both the lateral and sagittal derivations and the greater regional differentiation in lateral as compared to the sagittal channels. Development and aging over the course of the lifetime is characterized by considerable decreases in power from age 12 to age 20 in all frequency bands and in both lateral and sagittal orientations. In this age range the developmental pattern of trajectories of theta and alpha bands is quite dissimilar, with faster decreases in theta bands than in alpha bands, as well as frequency specific regional differences in the distribution of power values. The variation in power across scalp locations was very small in low theta and very large in high alpha. The variation in power in the theta band was primarily between central scalp locations and peripheral locations, while the variation in the alpha band was primarily between anterior and posterior locations. From the perspective of proportional change from the initial value, we find that by the age of 30, all low theta values were less than 90% of their values at age 12, while no high alpha values were less than 90% of their values at age 12; the majority of the central and posterior electrodes had values greater than 95% of their initial values (see Figures 7 through 10). In this period values for females are generally larger than values for males, and the rate of change (decrease) is larger in males than in females. Subsequently there are a variety of frequency and region dependent trajectories ranging from continued decreases in power in the theta bands to relative stability to increases, particularly in low alpha and in the beta frequency bands. Male power values were consistently lower than female values, but the shapes of the trajectories were similar from ages 12 to the early twenties, except in the high alpha band. Compared to the monopolar power, the bipolar power has consistently lower values and greatly reduced regional differences in the sagittal channels because of their topographical pattern. The lateral channels do not have the consistent increase in power from anterior to posterior of the monopolar power, but considerable overlap between central and posterior trajectories, possibly a result of greater phase synchrony in the posterior region. Beta power for bipolar channels increases much more after age 20 than it does for monopolar electrodes. This suggests that global phase synchrony in the beta frequency bands may be relatively high in the early development process and decreases considerably with age.

#### Quantification of sex differences

Using the relative Frobenius distance (RFD) method, male/female differences were calculated for each frequency band separately for all sagittal channels and all lateral channels. All values were less than 0.055, with the lateral distances (mean = 0.0428) slightly greater than the sagittal distances (mean = 0.0383). The differences were calculated over the entire lifespan; age range specific differences were not calculated, but might show considerable variation across ages. See table 1 for specifics.

**Table 1:**
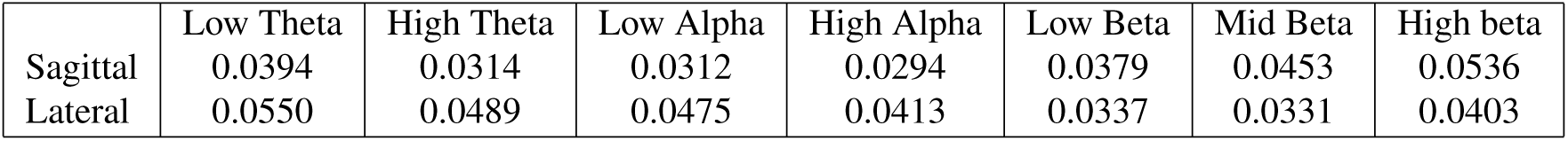
Male-Female RFD for power trajectories

### 3.3 Inter-Frequency correlation trajectories

The pairwise inter-frequency (within channel) correlation trajectories between the four theta and alpha frequency bands and the four high alpha and beta frequency bands at each channel show striking male-female differences unlike the corresponding monopolar correlations in Chorlian et al. (2026). Another phenomenon of interest is the presence of similar patterns of local extrema in the trajectories. Local minima in correlation trajectories suggest a relatively slow transition from one organization to another in interfrequency relations while local maxima suggest a rapid period of simplification followed by a more complex order. Exactly how this happens can only be understood by a comprehensive analysis of the trajectories of the relevant covariance matrices, as outlined in Chorlian et al. (2024).

#### 3.3.1 Theta-Alpha correlation trajectories

Inter-frequency theta-alpha correlations (shown in Figures 11 through 14) exhibit large sex differences. Practically all female trajectories have minima at age 40, more uniformly so for the lateral channels than the sagittal channels, which have some amount of fluctuation between ages 20 and 40. Male lateral trajectories have some minima before age 40, particularly for the central channels. but display less uniformity between regions than do the female trajectories. Male sagittal trajectories have local extrema, both maxima and minima, between ages 20 and 35, with subsequent variation, far less uniform than the female trajectories. The arrangement of the plots places the adjacent frequency correlations on the left and the more distant correlations on the right, with the design of having the overall level of correlation decrease from left to right. This occurs, but additionally there is a wider separation between the regions from left to right more extreme in the lateral channels than in the sagittal channels.

**Figure 7:**
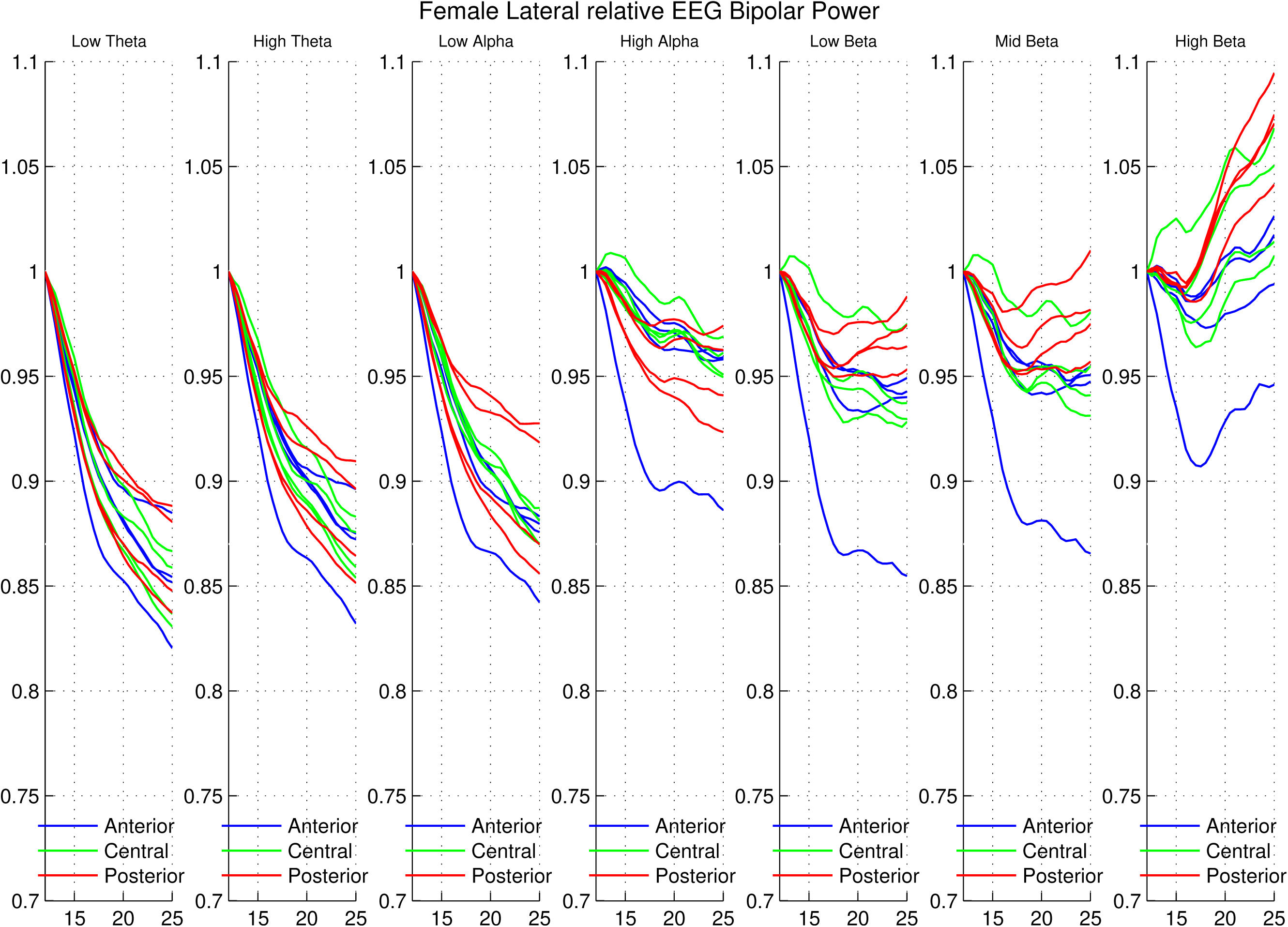
Relative power of the seven frequency band lateral female means to age 25

**Figure 8:**
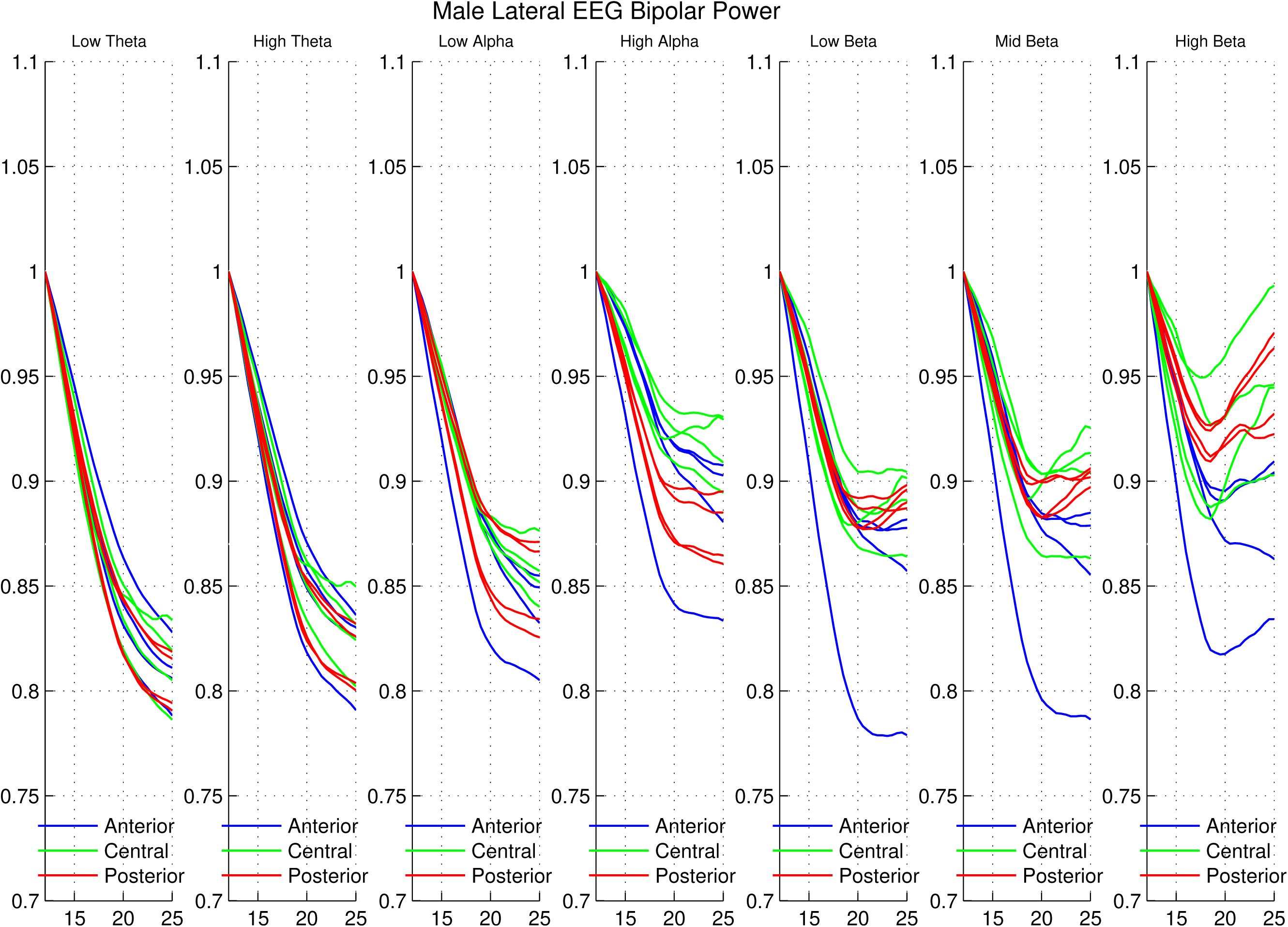
Relative power of the seven frequency band lateral male means to age 25

**Figure 9:**
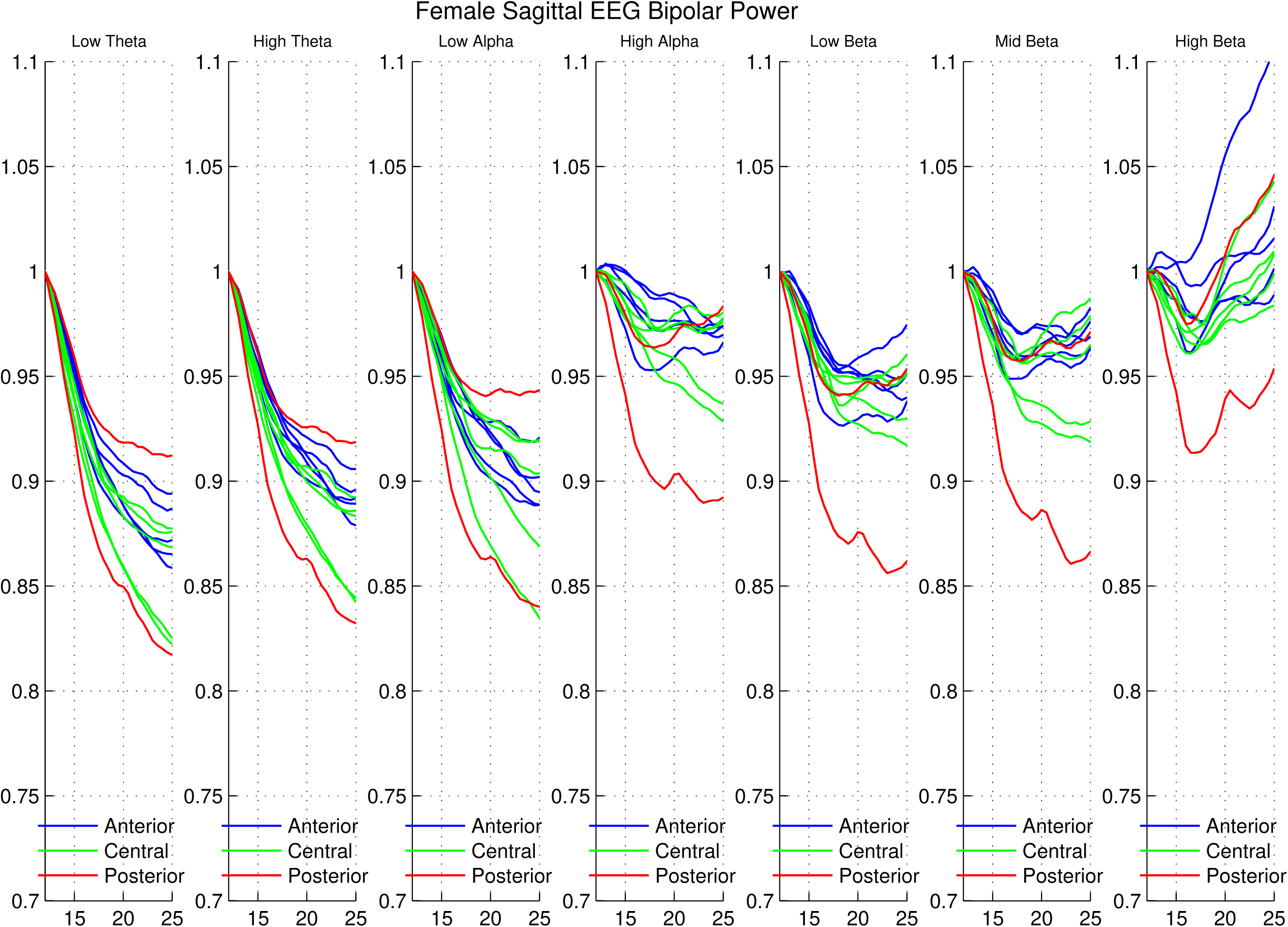
Relative power of the seven frequency band lateral female means to age 25

**Figure 10:**
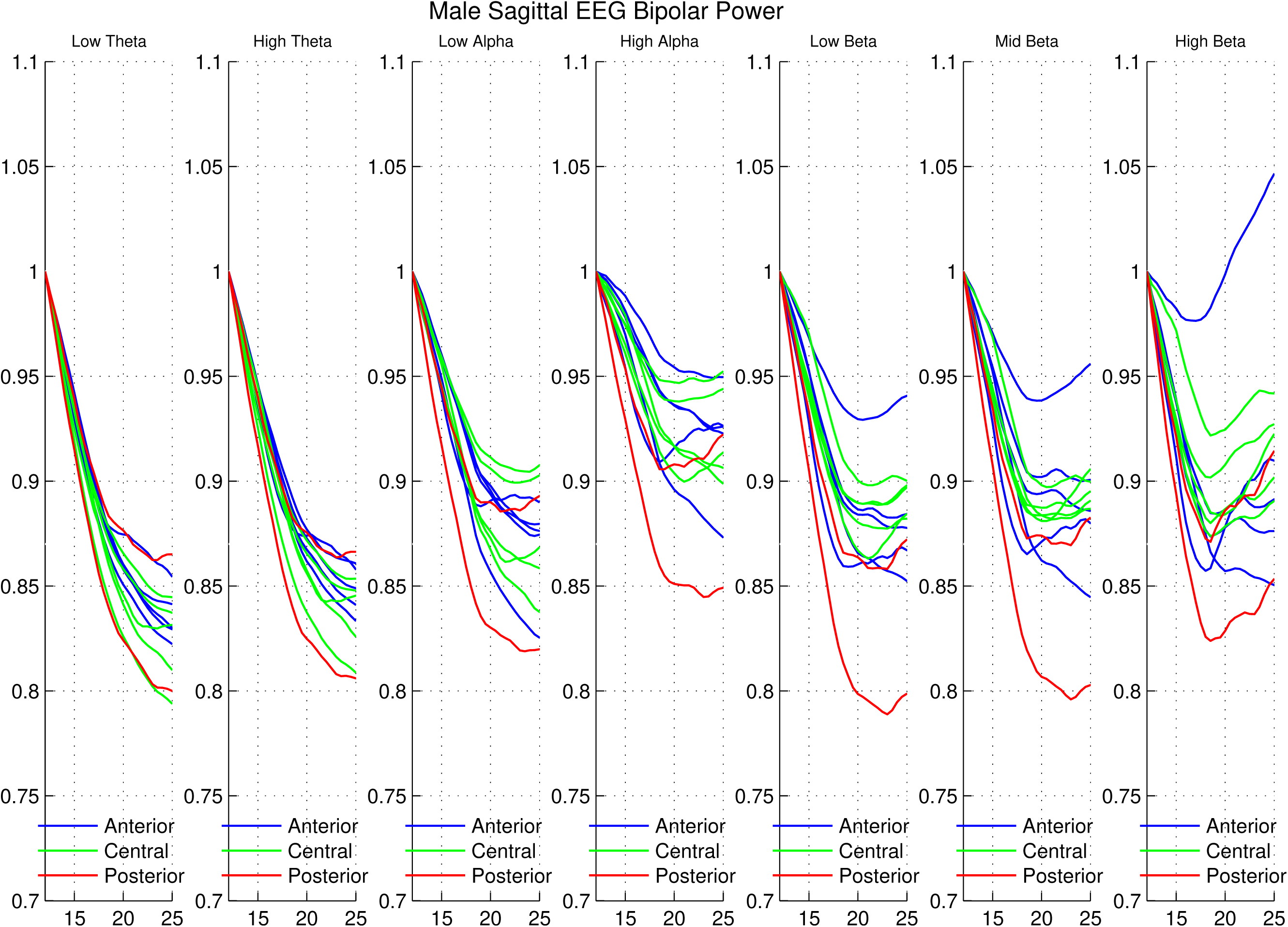
Relative power of the seven frequency band sagittal male means to age 25

**Figure 11:**
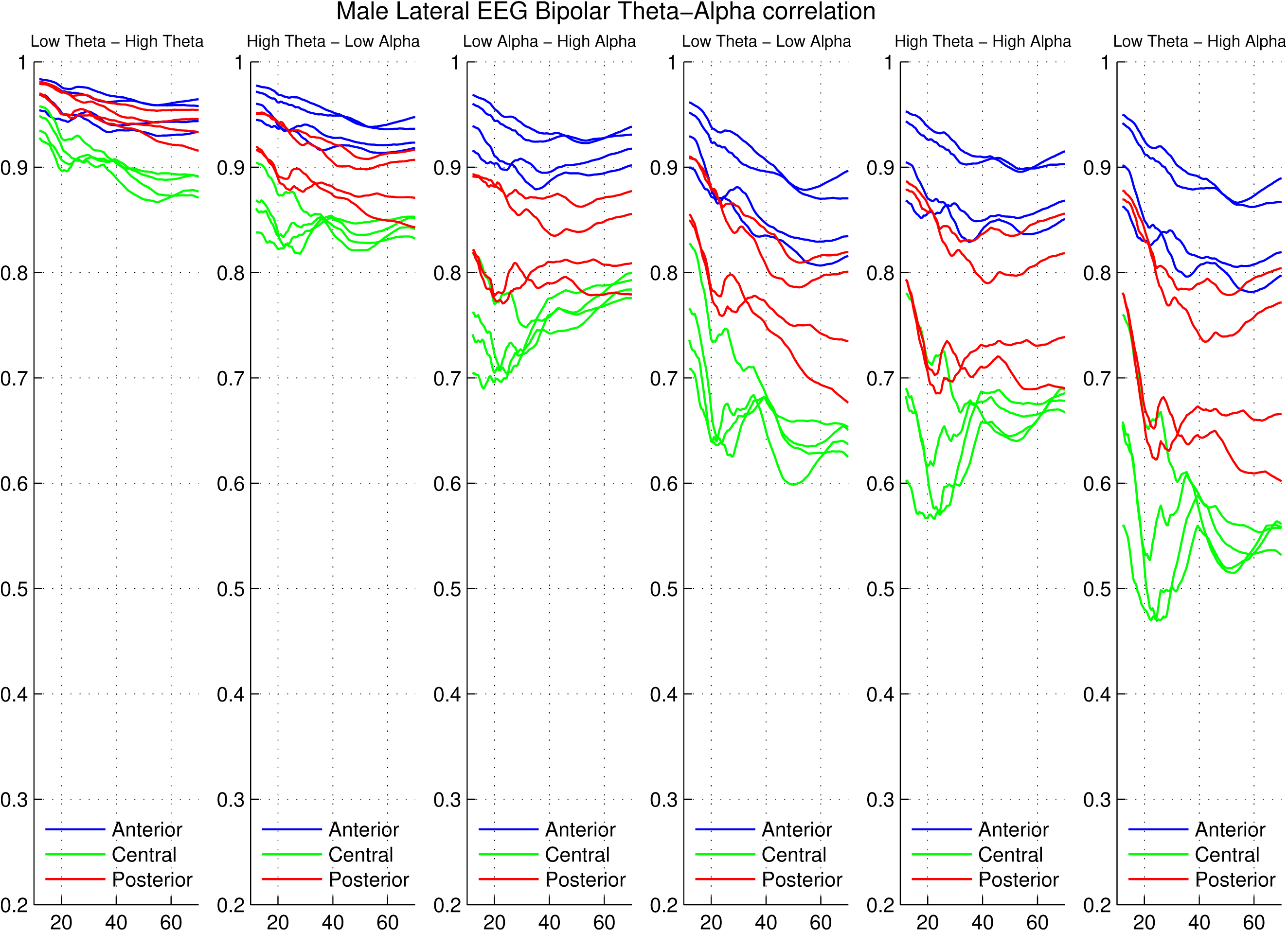
Trajectories of the six inter-frequency male lateral theta-alpha correlation pairs.

#### 3.3.2 Alpha-Beta correlation trajectories

Inter-frequency alpha-beta correlations (shown in Figures 15 through 18) have many of the characteristics of the thetaalpha correlations, except that the use of high alpha to provide a six column plot similar to the theta-alpha plots introduces a somewhat anomalous element to the within beta correlations which are much higher than the beta-high alpha correlations.

**Figure 12:**
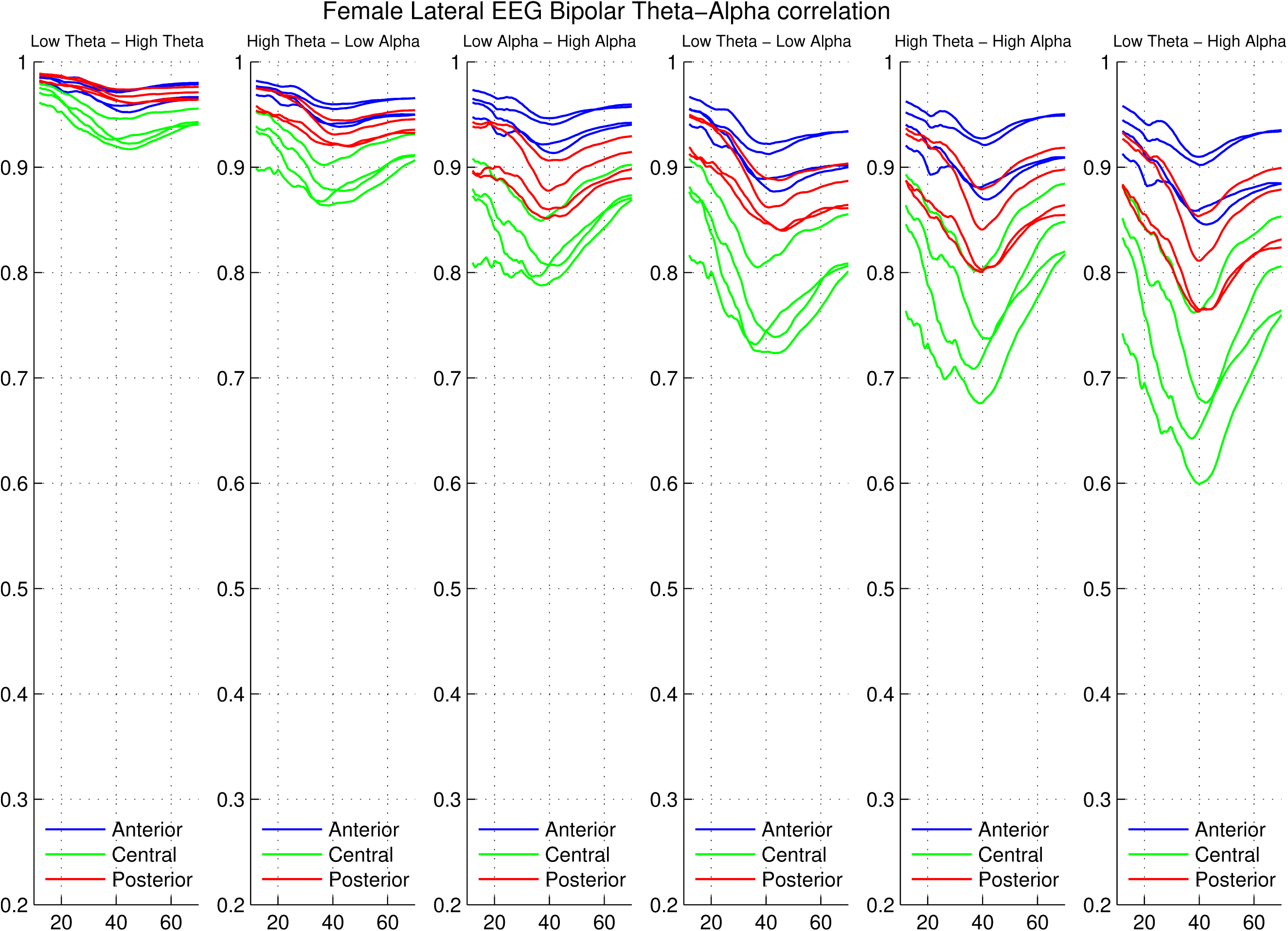
Trajectories of the six inter-frequency female lateral theta-alpha correlation pairs

**Figure 13:**
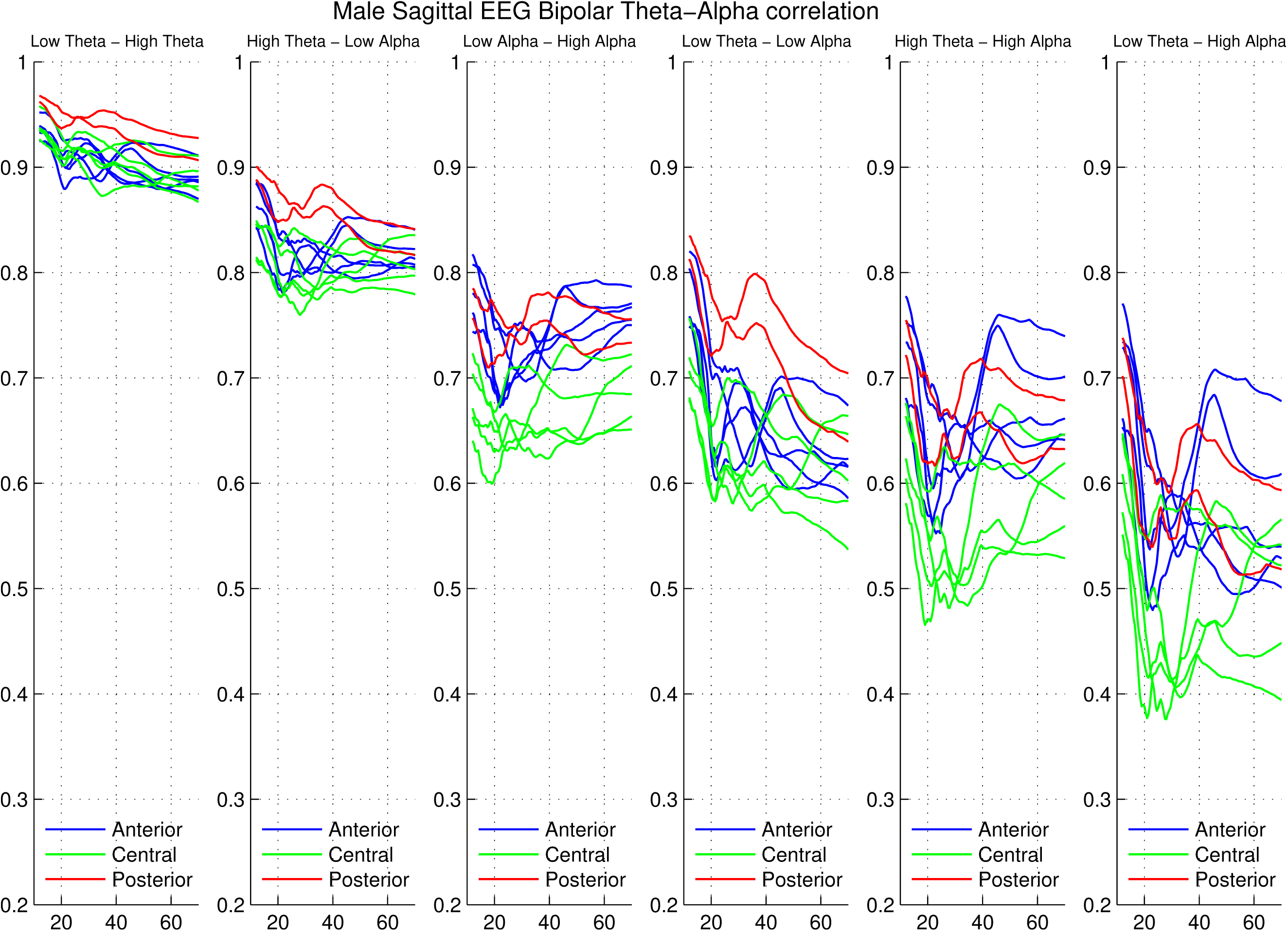
Trajectories of the six inter-frequency male sagittal theta-alpha correlation pairs

**Figure 14:**
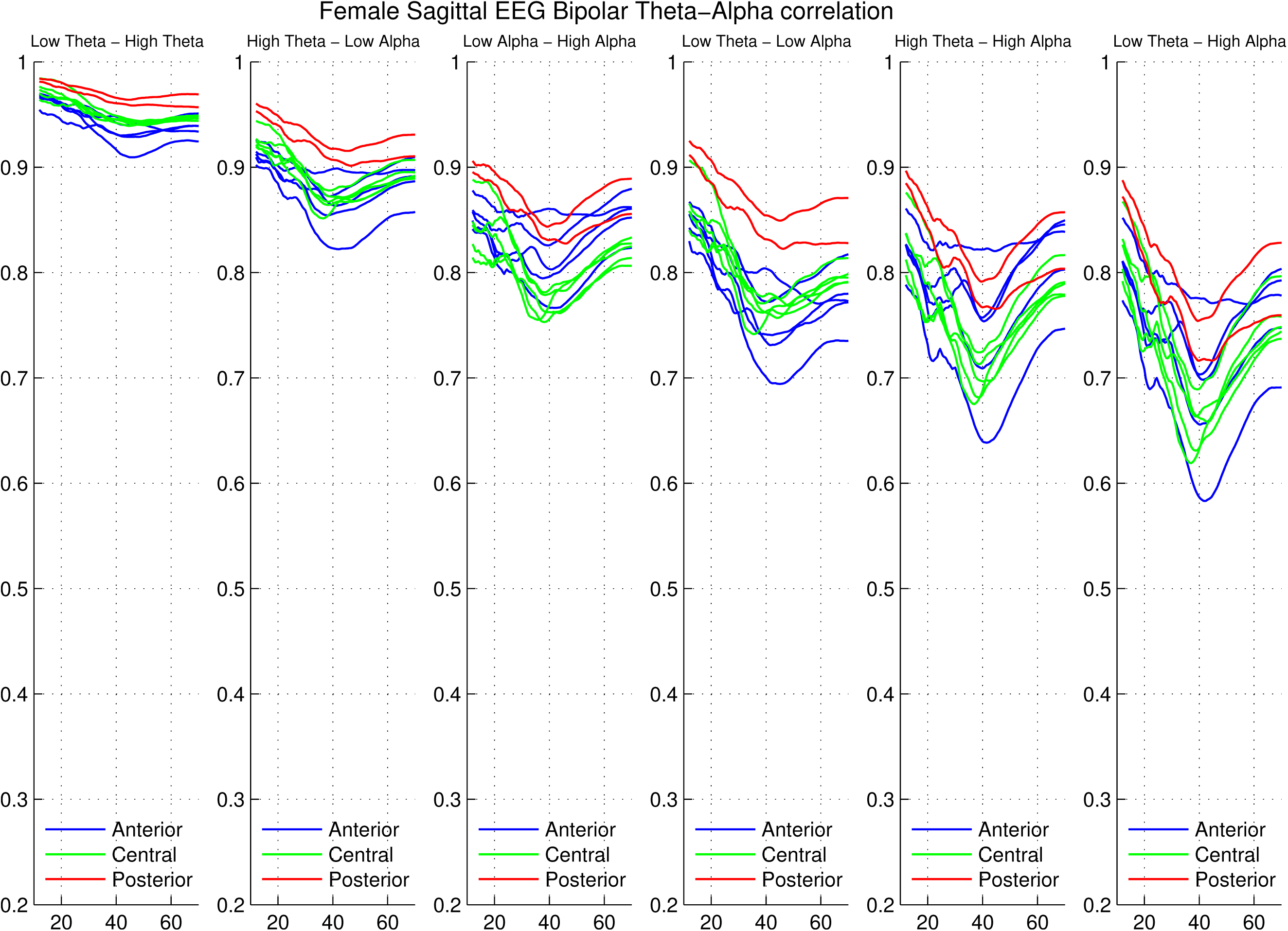
Trajectories of the six inter-frequency female sagittal theta-alpha correlation pairs

**Figure 15:**
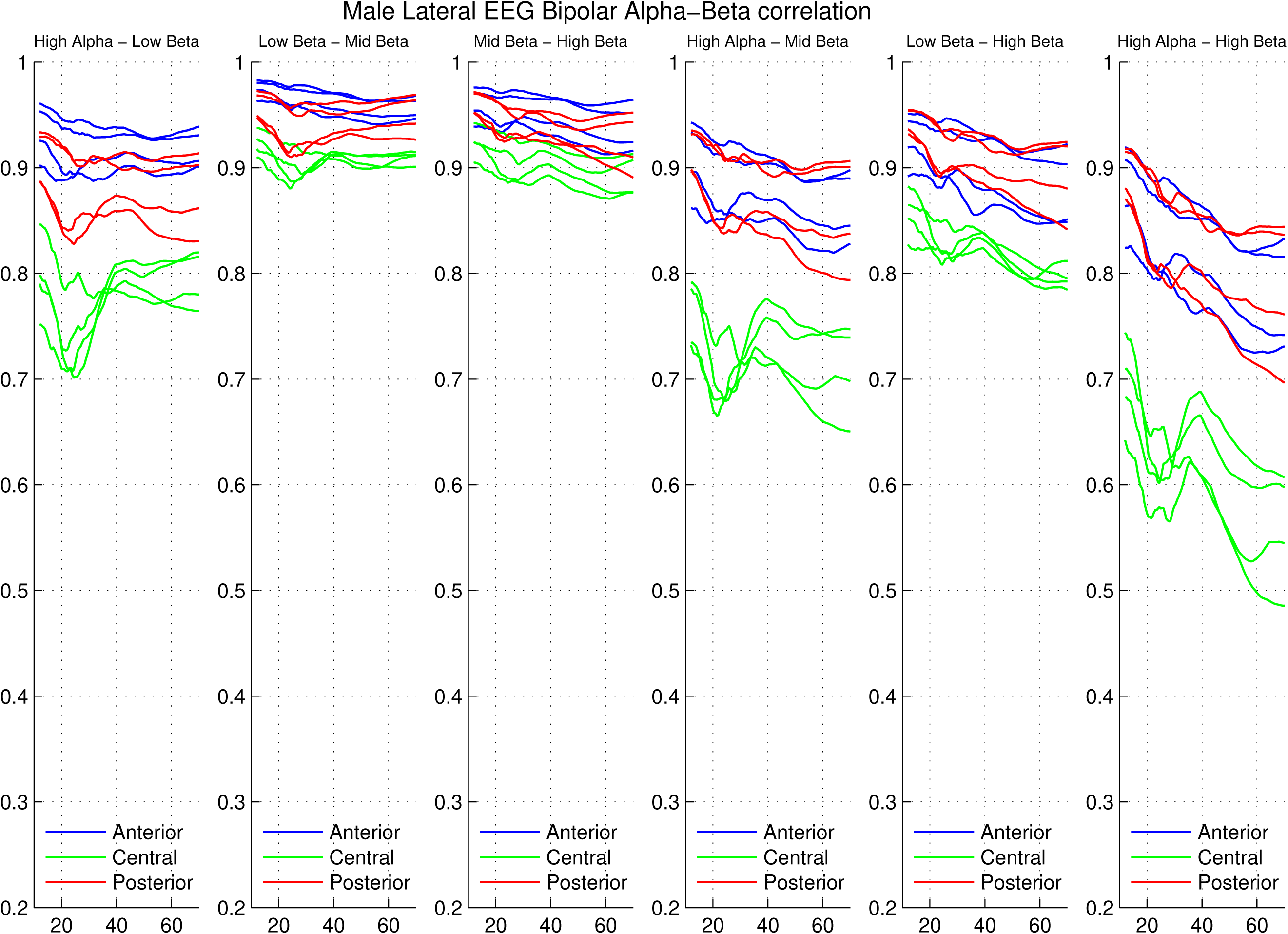
Trajectories of the six inter-frequency male lateral alpha-beta correlation pairs.

##### Quantification of sex differences

Using the relative Frobenius distance method, male/female differences were calculated for each of the six inter-frequency pairs in each of the two channel groups for the theta-alpha group and the alpha-beta group of frequencies. Theta-alpha sagittal RFDs ranged from .047 to .318 and lateral values from .034 to .177 with wider spaced between frequency comparisons having larger values. Alpha-beta sagittal RFDs ranged from .042 to .169 and lateral values from .026 to .119 between with wider spaced between frequency comparisons having larger values. Theta-alpha values were always larger than Alpha-beta values except for LT-HT and HT-LA correlations. These values are considerably larger than those for the power trajectories. See tables 2 and 3 for specifics.

**Table 2:** Male-Female RFD for Theta-Alpha correlation trajectories

|  | LT-HT | HT-LA | LA-HA | LT-LA | HT-HA | LT-HA |
| --- | --- | --- | --- | --- | --- | --- |
| Sagittal | 0.0467 | 0.0892 | 0.1558 | 0.2057 | 0.2344 | 0.3176 |
| Lateral | 0.0337 | 0.0502 | 0.0799 | 0.1222 | 0.1236 | 0.1769 |

**Table 3:** Male-Female RFD for Alpha-Beta correlation trajectories

|  | HA-LB | LB-MB | MB-HB | HA-MB | LB-HB | HA-HB |
| --- | --- | --- | --- | --- | --- | --- |
| Sagittal | 0.1079 | 0.0417 | 0.0395 | 0.1063 | 0.0705 | 0.1692 |
| Lateral | 0.0656 | 0.0288 | 0.0263 | 0.0780 | 0.0531 | 0.1194 |

### 3.4 Intra-Frequency correlation trajectories

The pair-wise inter-channel correlations for the 7 frequency bands, examined for 23 inter-channel pairs, show little between frequency difference within each pair. Since we do not have a satisfactory method for quantifying the similarity, there will be no further discussion of them.

### 3.5 Comparison to monopolar results

#### 3.5.1 Similarity of power trajectories

The features mentioned above about the similarity between the bipolar trajectories, particularly the lateral trajectories, and the monopolar trajectories are clear on examination of the plots of the monopolar trajectories, as shown in Figures 1 and 2 of Chorlian et al. (2026) and reproduced here as Figures 19 and 20.

**Figure 16:**
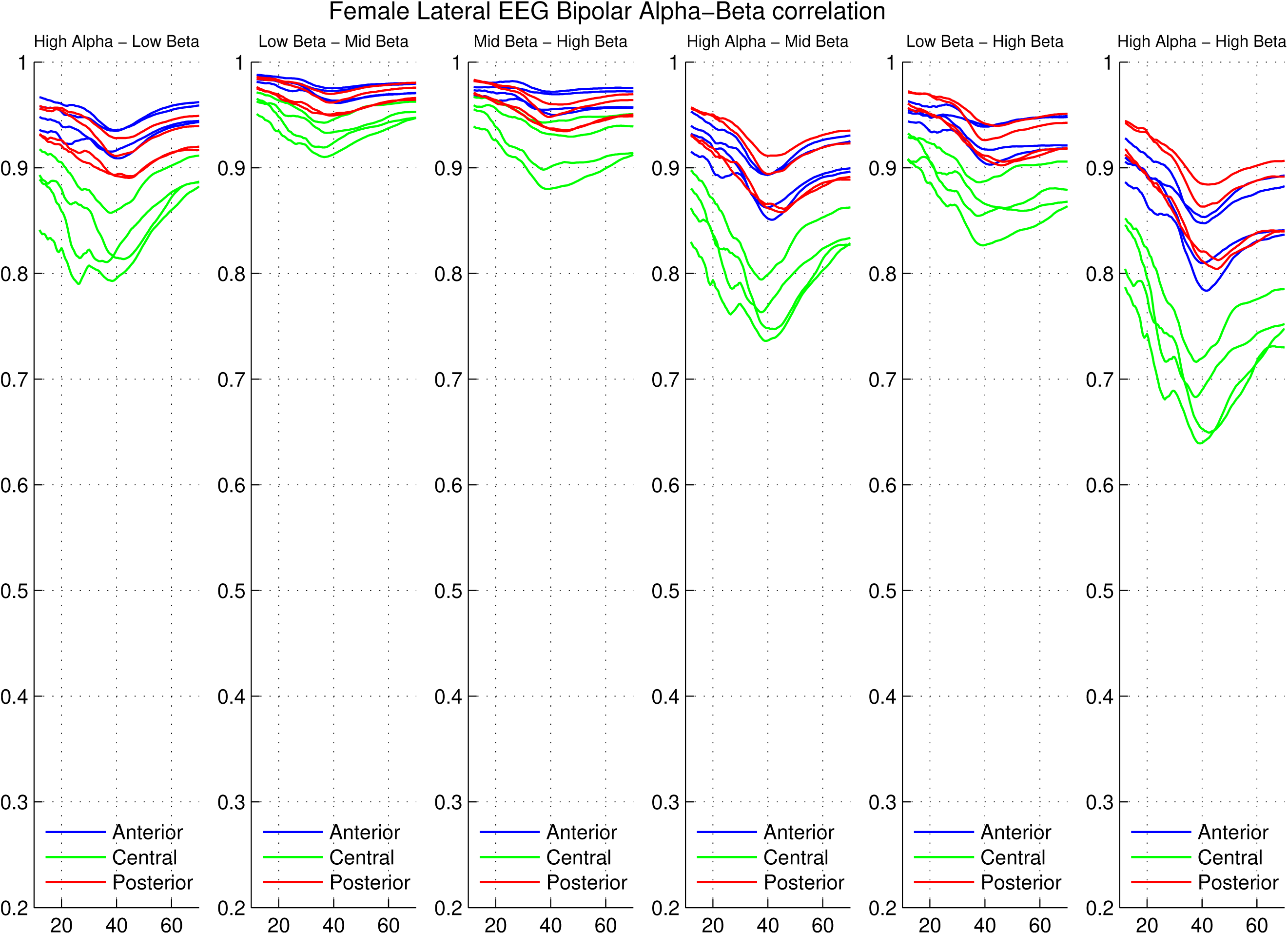
Trajectories of the six inter-frequency female lateral alpha-beta correlation pairs

**Figure 17:**
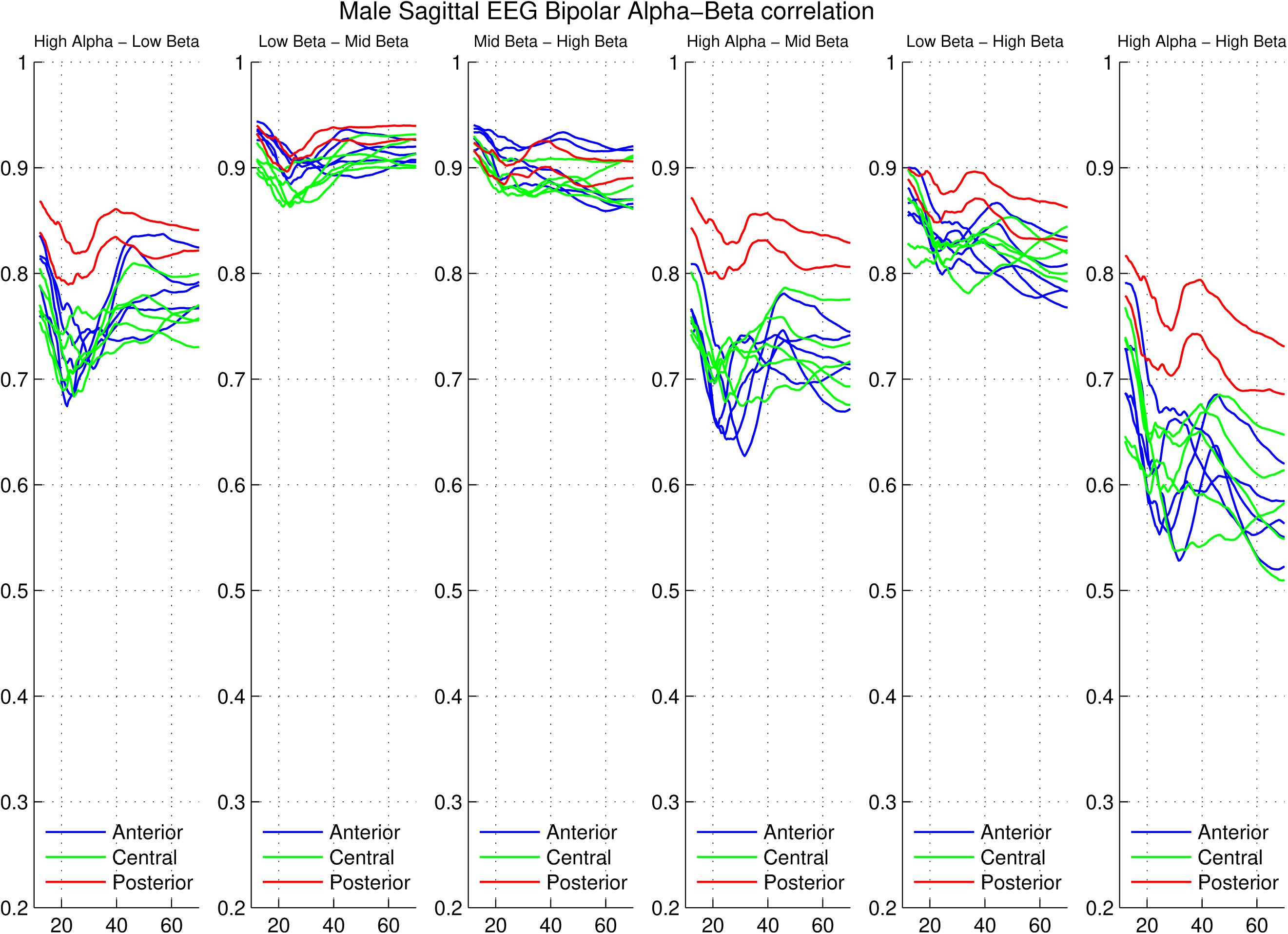
Trajectories of the six inter-frequency male sagittal alpha-beta correlation pairs

**Figure 18:**
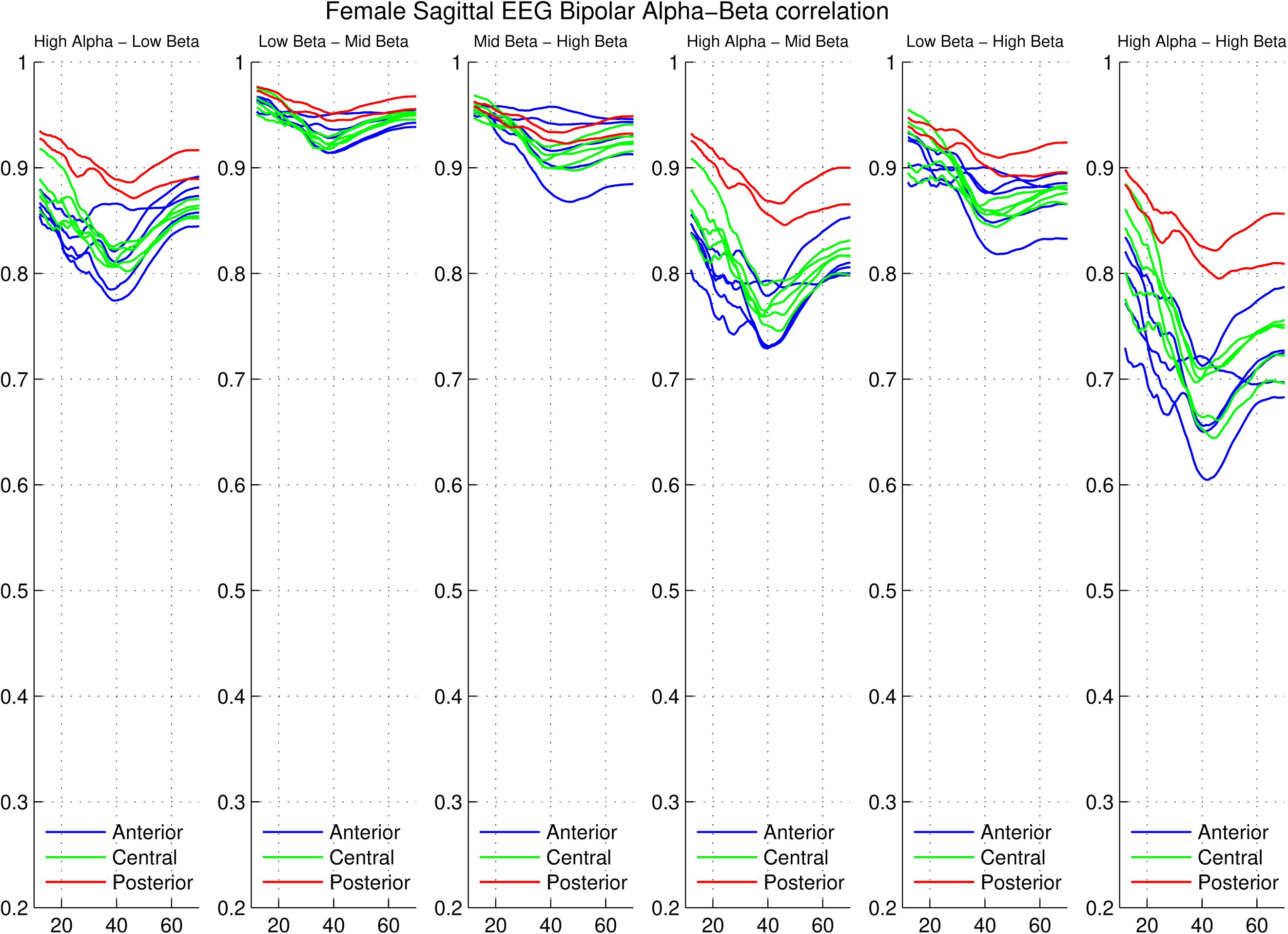
Trajectories of the six inter-frequency female sagittal alpha-beta correlation pairs

**Figure 19:**
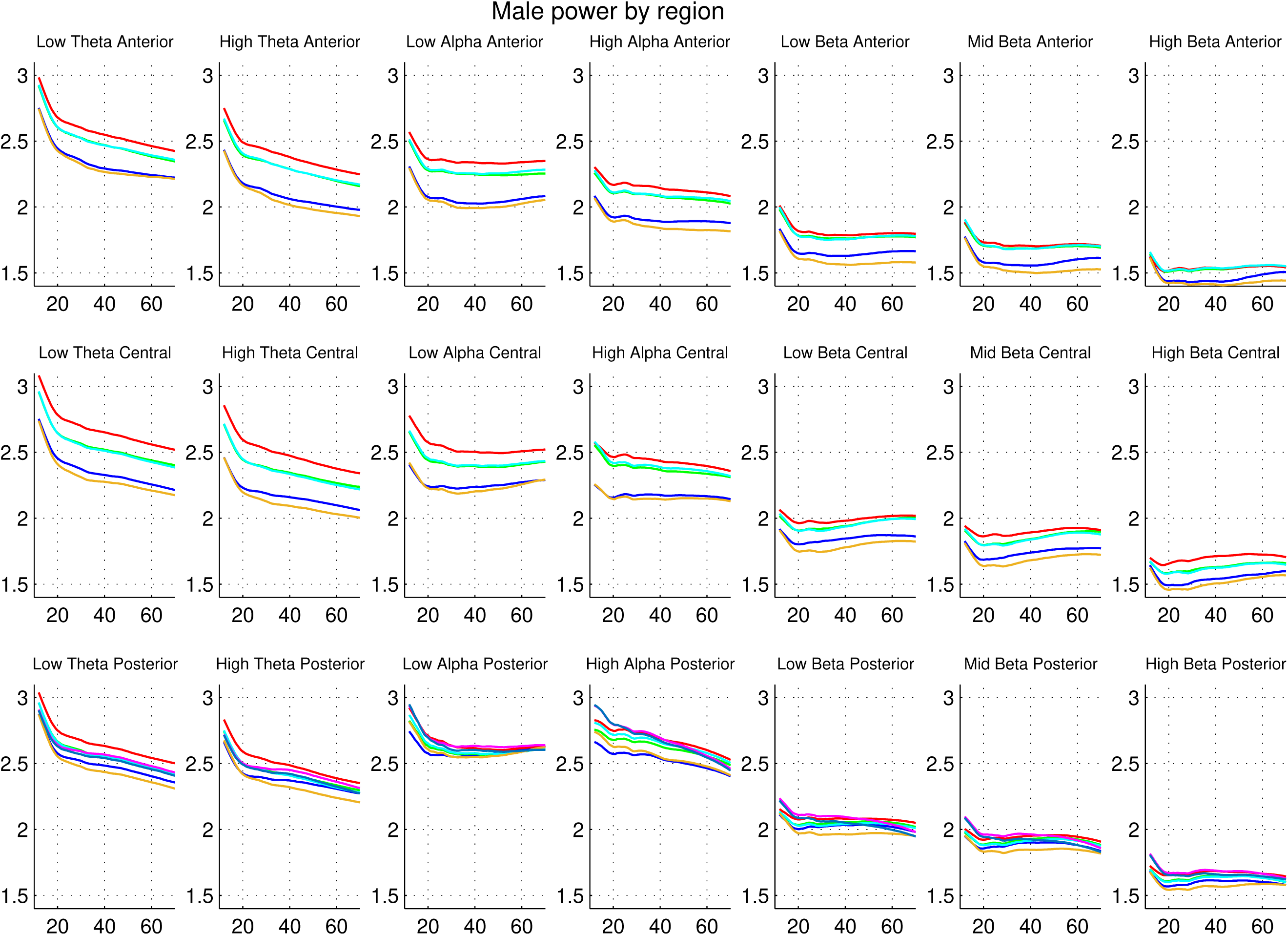
Monopolar male trajectories. Color coding from left to right is blue, green, red, cyan, mustard.

**Figure 20:**
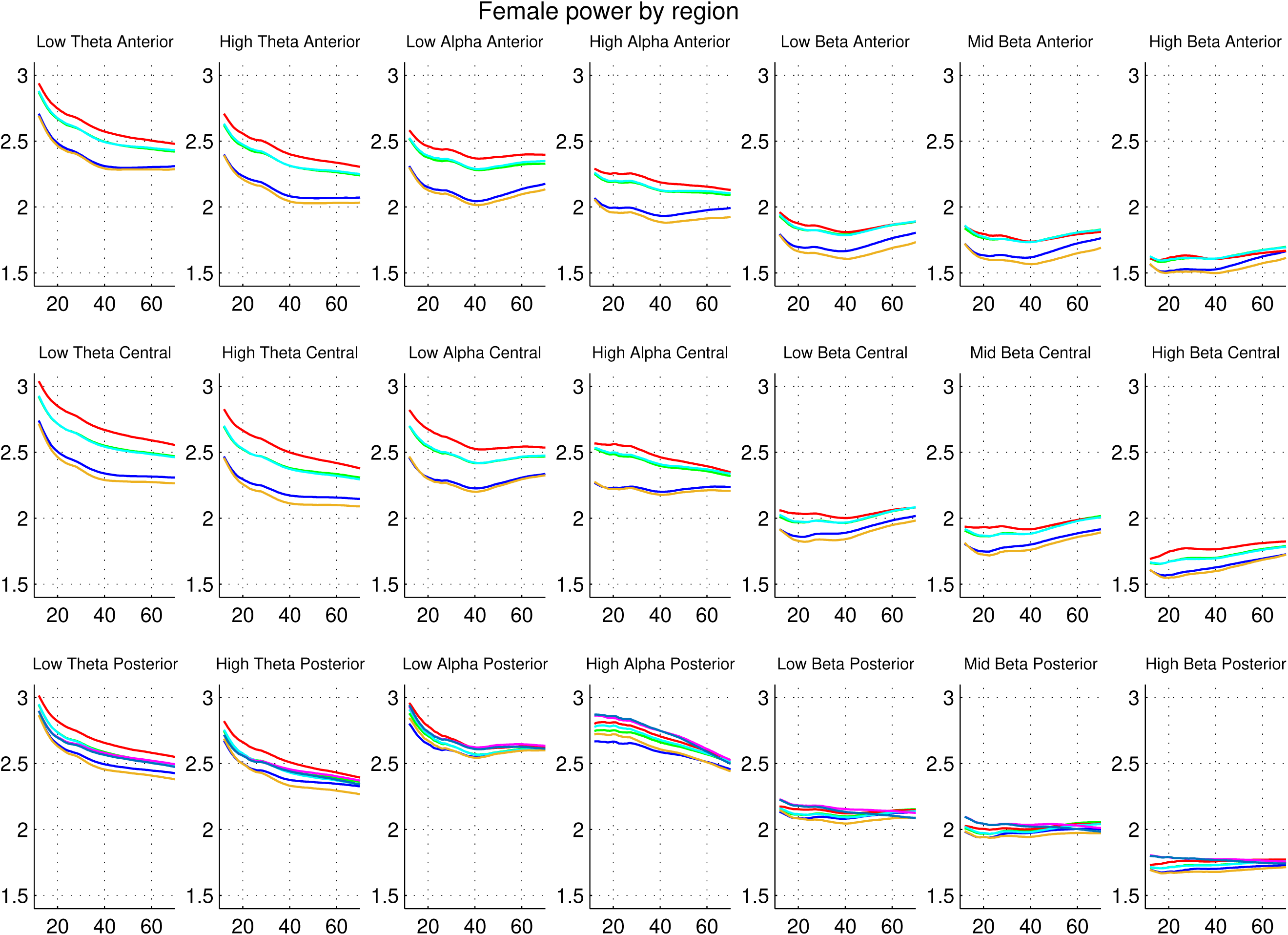
Monopolar female trajectories. Color coding from left to right is blue, green, red, cyan, mustard.

#### 3.5.2 Effect of phase synchrony on power trajectories

As mentioned previously, there are a number of features in the bipolar power trajectories that suggest that the differencing procedure reduces not only local phase synchrony, but also effects of global and regional phase synchrony present in the monopolar data from which the bipolar signals are derived. In the lateral bipolar channels, the peripheral channels have greater power than the internal channels, exactly the opposite of the case in the monopolar channels, as a consequence of the assumed greater central effect of larger scale synchrony. This is illustrated in figures 19 and 20 based on data from our monopolar study illustrating the monopolar channels and 21 and 22 illustrating the comparable lateral bipolar channels. Since there are 5 monopolar channels per region and only 4 bipolar channels, the color matching of comparable pairs is not exact, but the left peripheral channel is blue in each panel, and the right peripheral channel is cyan in the bipolar panels, and yellow in the monopolar channels. The interior channels are red and green in each panel and in the monopolar panels also cyan, which has in most cases overwritten the green although there are small patches where both are simultaneously visible.

#### 3.5.3 Dissimilarity of correlation trajectories

As mentioned previously the levels of inter-frequency correlation are for the bipolar channels are generally larger and in many cases much larger than those for the monopolar channels, as can be seen by comparing Figures 11 through 18 with Figures 23 and 24 taken from Chorlian et al. (2026). Since the phase synchrony has been reduced in the bipolar data, this suggests that global and regional phase synchrony components have greater frequency specificity than the more purely local components. This is consistent with the apparent lack of frequency specificity in the inter-channel correlation trajectories mentioned above.

**Figure 21:**
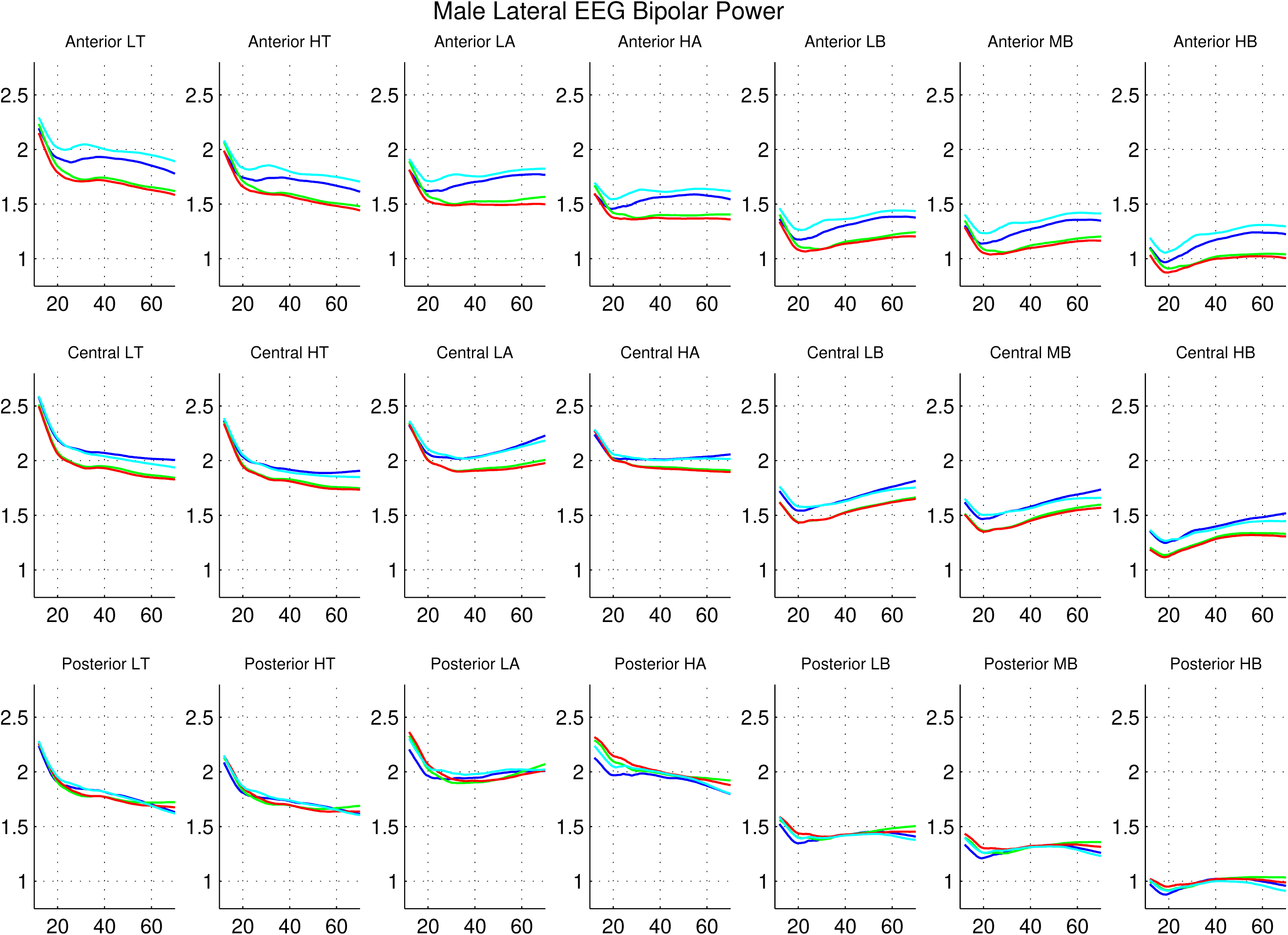
Lateral (bipolar) male trajectories. Color coding from left to right is blue, green, red, cyan.

**Figure 22:**
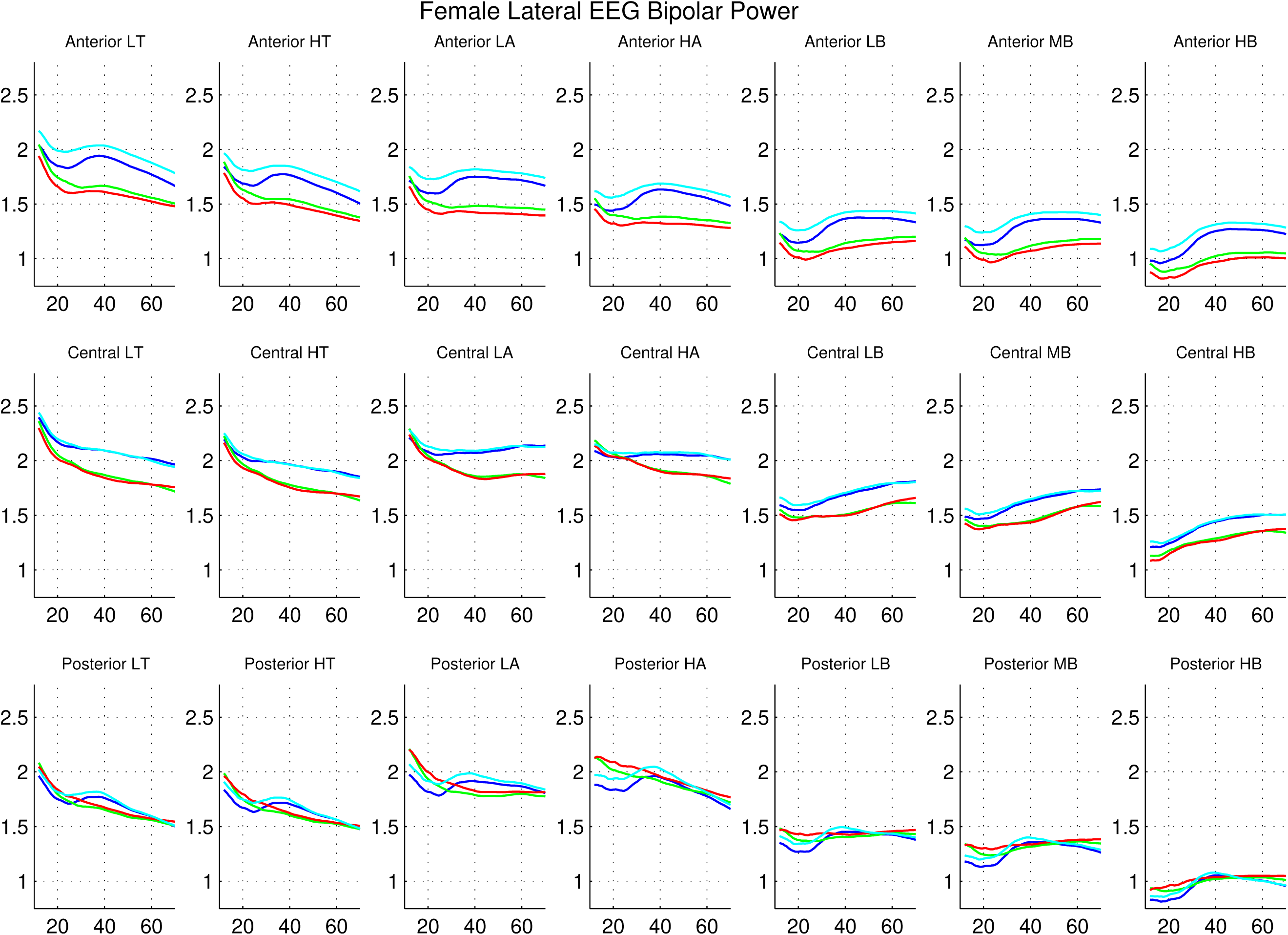
Lateral (bipolar) female trajectories. Color coding from left to right is blue, green, red, cyan.

**Figure 23:**
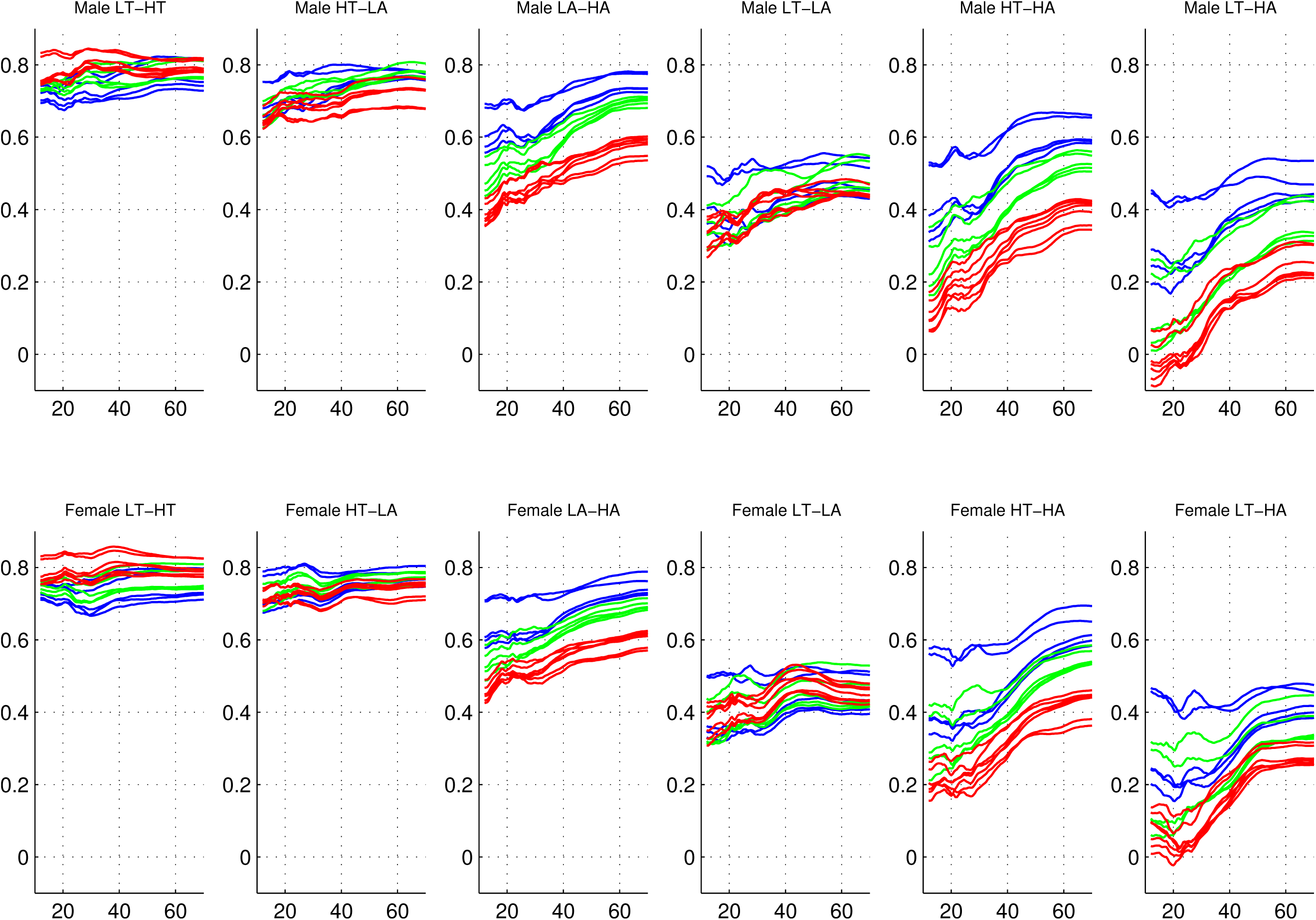
Trajectories of the male and female theta-alpha band inter-frequency correlations of monopolar data. Electrodes are color coded by region.

**Figure 24:**
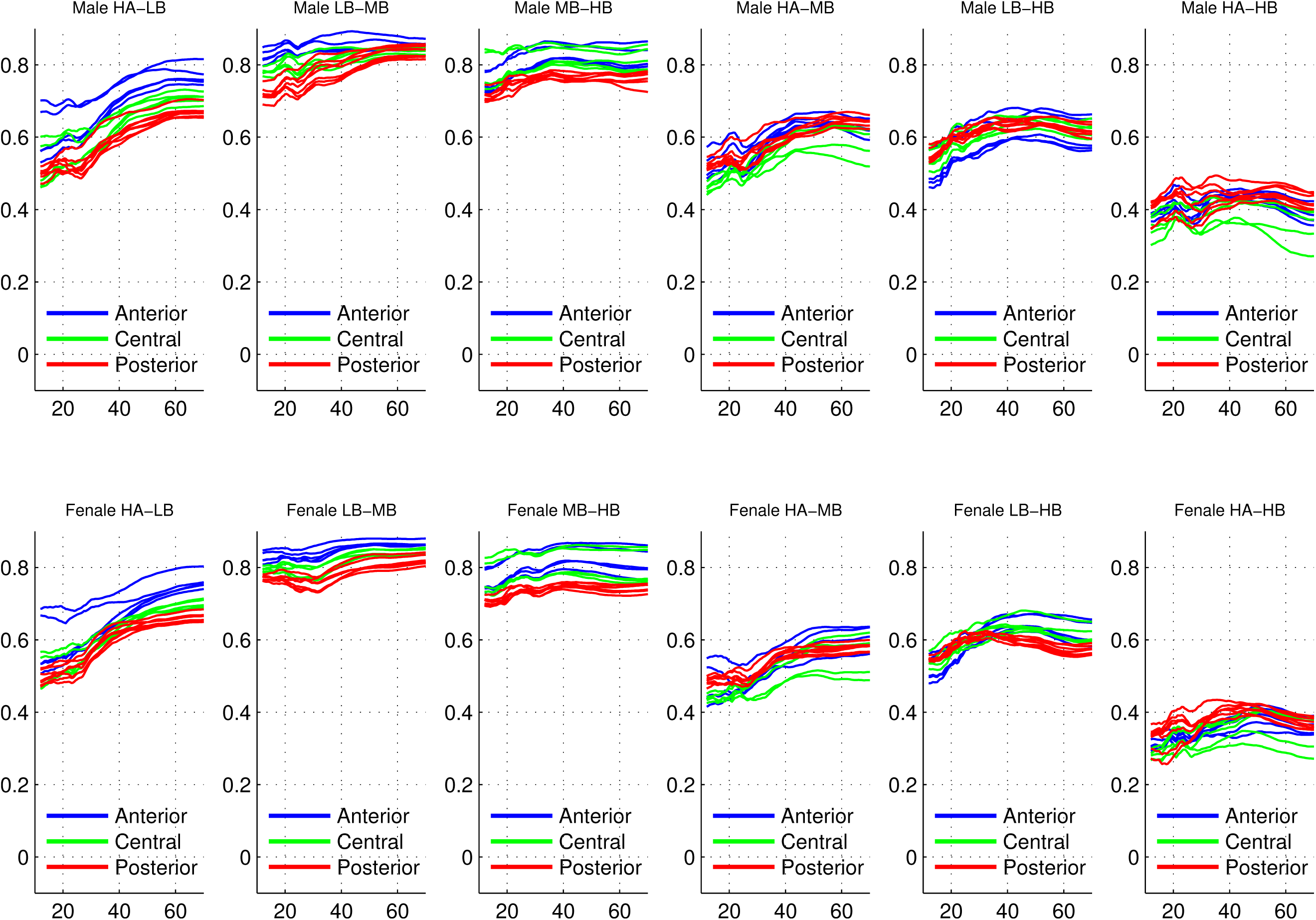
Trajectories of the male and female alpha-beta band inter-frequency correlations of monopolar data. Electrodes are color coded by region.

## 4 Extensions of this analysis

As mentioned previously, the RFD method used to quantify male/female differences in individual frequency and channel groups could not be used to directly compare sagittal and lateral channels as two distinct groups. However, the examination of aggregated results from separate computations such as male/female differences in the trajectories of covariance structure of lateral and sagittal channels could be informative about differences between males and females in the spatial organization of brain function. Refinement of our methods of comparing trajectories would enable quantification of the overall similarity of trajectories across a range of measures, particularly for the inter-channel correlation trajectories.

Thus the next stage of analysis is envisioned as the direct analysis of the covariance matrices,to provide trajectories of larger aggregates, such as encompassing all frequency bands for individual channels or all appropriate channel for individual frequency bands, based on their pair-wise distances as described and used in Chorlian et al. (2024). Using the same methodology on the monopolar data would provide an additional layer of analysis to understand the effect of the differencing process.

## 5 Electrode, Channel, and Frequency Structure

### 5.1 Frequency bands

Frequency bands:

1. Low theta 3.0-5.0 Hz. 2. High theta 5.0-7.0 Hz. 3. Low alpha 7.0-9.0 Hz. 4. High alpha 9.0-12.0 Hz. 5. Low beta 12.0-16.0 Hz.
2. High beta 16.0-20.0 Hz. 7. High beta 20.0-28.0 Hz.

### 5.2 Regional structure of Electrodes

Frontal:

1. F7; 2. F3; 3. FZ; 4. F4; 5. F8;

Central:

1. T7; 7. C3; 8. CZ; 9. C4; 10. T8;

Parietal:

1. P7; 12. P3; 13. PZ; 14. P4; 15. P8;

Occipital:

1. O1; 17. O2;

### 5.3 Electrode structure of Channels

Sagittal frontal:

1. F7-T7; 2 F3-C3; 3 FZ-CZ; 4 F4-C4; 5 F8-T8;

Sagittal central:

1. T7-P7; 7 C3-P3; 8 CZ-PZ; 9 C4-P4; 10 T8-P8;

Sagittal posterior:

1. P3-O1; 12 P4-O2;

Lateral frontal:

1. F7-F3 14 F3-FZ 15 FZ-F4 16 F8-F4

Lateral central:

1. T7-C3 18 CZ-C3 19 CZ-C4 20 T8-C4

Lateral parietal:

1. P7-P3 22 PZ-P3 23 PZ-P4 24 P8-P4

## 6 Figures

The initial comprehensive views provide the trajectories of the power values across the entire age range, the entire set of frequency bands, and the entire set of channels for males and female in separate plots. Following these are inter-frequency correlation plots, with frequency bands labeled by pairs of letters, i.e, "LT" = low theta etc. Male and female values are on separate plots.

## 8 Acknowledgments

The Collaborative Study on the Genetics of Alcoholism (COGA), Principal Investigators B. Porjesz, V. Hesselbrock, A. Agrawal; Scientific Director, A. Agrawal; Translational Director, D. Dick, includes nine different centers: University of Connecticut (V. Hesselbrock); Indiana University (H.J. Edenberg, T. Foroud, Y. Liu, M.H. Plawecki); University of Iowa Carver College of Medicine (A. Andersen S. Kuperman); SUNY Downstate Health Sciences University (B. Porjesz, J. Meyers); Washington University in St. Louis (L. Bierut, A. Agrawal, S. Hartz); University of California at San Diego (M. Schuckit); Rutgers University (D. Dick, R. Hart, J. Salvatore, J. Tischfield); The Children’s Hospital of Philadelphia, University of Pennsylvania (L. Almasy); Icahn School of Medicine at Mount Sinai (A. Goate, P. Slesinger); and Howard University (D. Scott). Other COGA collaborators include: M. Hesselbrock, K. Manning (University of Connecticut); D. Lai, J. Nurnberger Jr., L. Wetherill, A. Miller, X. Xuei, (Indiana University); J. Kramer (University of Iowa), G. Chan (University of Iowa; University of Connecticut); C. Kamarajan, A. Pandey, D.B. Chorlian, P. Barr, S. Kinreich, G. Pandey, Z. Neale, C. Chatzinakos, J. Zhang, S. Saenz deViteri, A. Bingly (SUNY Downstate); G. Pathak (Icahn School of Medicine at Mount Sinai); A. Anokhin, K. Bucholz, F. Dong, A. Hatoum, E. Johnson, J. Rice, S. Saccone (Washington University); F. Aliev, Z. Pang, S. Kuo, S. Brislin, (Rutgers University). We continue to be inspired by our memories of Henri Begleiter and Theodore Reich, founding PI and Co-PI of COGA, and also owe a debt of gratitude to other past organizers of COGA, including Ting-Kai Li, P. Michael Conneally, Raymond Crowe, and Wendy Reich, for their critical contributions. This national collaborative study is supported by NIH Grant U10AA008401 from the National Institute on Alcohol Abuse and Alcoholism (NIAAA) and the National Institute on Drug Abuse (NIDA).

## Notes

### Competing Interest Statement

The authors have declared no competing interest.

